# The multidimensional nutritional niche of fungus-cultivar provisioning in free-ranging colonies of a neotropical leafcutter ant

**DOI:** 10.1101/2021.02.22.432261

**Authors:** Antonin J.J. Crumière, Aidan James, Pol Lannes, Sophie Mallett, Anders Michelsen, Riikka Rinnan, Jonathan Z. Shik

## Abstract

The foraging trails of *Atta* leafcutter colonies are among the most iconic scenes in Neotropical ecosystems, with thousands of ants carrying freshly cut plant fragments back to their nests where they are used to provision a fungal food crop. We tested a hypothesis that the fungal cultivar’s multidimensional requirements for macronutrients (protein and carbohydrates) and minerals (Al, Ca, Cu, Fe, K, Mg, Mn, Na, P and Zn) govern the foraging breadth of *Atta colombica* leafcutter ants in a Panamanian rainforest. Analyses of freshly cut plant fragments carried by leafcutter foragers showed that the combination of fruits, flowers, and leaves provide for a broad realized nutritional niche that can maximize cultivar’s performance. And, while the leaves that comprised the most harvested resource also delivered an intake target containing protein in excess of the amounts that can maximize cultivar growth, *in vitro* experiments showed that the minerals P, Al, and Fe can enhance the cultivar’s tolerance to protein-biased substrates, and potentially expand the ants’ foraging niche. Yet, the cultivar also exhibits narrow margins between mineral limitation and toxicity that may render plant fragments with seemingly optimal blends of macronutrients unsuitable for provisioning. Our approach highlights that optimal foraging is inherently multidimensional and links the foraging behavior of a generalist insect herbivore to the fundamental nutritional niche of its microbial symbiont.

**Significance Statement:** Colonies of *Atta colombica* leafcutter ants can contain millions of specialized workers exhibiting large-scale generalist herbivory. Yet, this generalist foraging niche also depends on the poorly understood physiological needs of the ants’ domesticated fungal cultivar. We show the cultivar’s fundamental nutritional niche is broad for carbohydrates but narrower for protein and a suite of minerals, but that the cultivar’s sensitivity to excess protein is also mediated by Al, Fe, and P. More generally, this study decouples the multidimensional foraging strategies that enable a generalist herbivore to navigate a complex nutritional landscape and mix many imbalanced foods to achieve balanced cultivar provisioning.

## Main Text

### Introduction

Natural selection is predicted to favor traits that enable consumers to acquire nutritionally balanced diets (1, 2). For insect herbivores, such nutrient regulation often poses major challenges. First, plant foods tend to contain carbon (C) in far higher concentrations than other limiting resources like nitrogen (N) and phosphorus (P) (3). Second, each mouthful of ingested plant tissue is likely to contain valuable macronutrients (e.g. carbohydrates, proteins, lipids) and other essential components (e.g. vitamins, minerals), but also a mix of recalcitrant compounds (e.g. cellulose) and toxins (e.g. tannins) (4). Third, insects are seldom limited by a single nutrient at a time, and the value of a given plant resource thus depends on the ratios and concentrations of multiple interacting nutrients (5). The field of nutritional geometry has provided new approaches for studying these multidimensional dietary challenges (6-8) and has shown that organisms have diverse strategies for prioritizing specific nutrients when foraging for and consuming imbalanced foods (9-11). We extended nutritional geometry approaches to study nutritional regulation strategies in free-ranging colonies of the leafcutter ant *Atta colombica*. These ants are ecologically important neotropical herbivores and belong to a lineage that is unique among ants in collecting food resources (*i*.*e*. plant fragments) to provision a domesticated fungal food crop (*Leucoagaricus gongylophorus*) rather than ant nestmates (12).

Nutritional geometry studies have usually focused on covarying macronutrients ((13-15), *but see* (16)), even though over 25 mineral elements are essential for life (17, 18). For instance, leafcutter ants concentrate Mg and Ca in their cuticle as a protective armor (19) and Zn as a hardening agent in their mandibles (20), while also preferentially foraging for Na-rich substrates (21) and avoiding vegetation with elevated Mn and Al (22). Plant foods are typically assumed to contain minerals in sufficient abundance to meet the low requirements of insect herbivores (4), but mineral concentrations also vary widely across plant species and tissues within individual plants (23-25). Minerals also tend to exhibit thresholds beyond which limitation becomes toxicity (26, 27), and minerals like Na, Al, Fe, Cu, and Zn can even be sequestered by plants to deter herbivores as quantitative chemical defenses (28-31). We thus hypothesized that minerals in vegetation can inhibit farming performance when leafcutter ants provision them in excess of their fungal cultivar’s tolerances and requirements.

Leafcutter ants have multiple opportunities for such nutritional regulation. First, each type of plant substrate has a specific nutritional profile (Figure 1A), and colonies can target nutritional blends by foraging among leaves, fruits, flowers (32, 33) and across plant species (Figure 1B) (34). Indeed, a single *A. colombica* colony can forage up to 126 plant species (53 families) and up to 370 kg (dry mass) of plant substrates during an annual cycle (35). These ants are thus extreme generalist foragers compared to the majority of insect herbivores that consume a few (ca. 3) plant families (36). The next phase occurs when gardener ants within underground fungus cultivation chambers macerate vegetation fragments and add a mixture of enzyme-rich fecal droplets to promote fungal hyphal growth and production of nutrient-rich hyphal tips called gongylidia (packaged in bundles called staphylae) (Figure 1C) (37, 38). Given the potential fungicultural benefits of optimized nutrient provisioning, we conjecture that colonies forage across plant substrates (Figure 1D) to acquire a realized nutritional niche (RNN) that targets cultivar fundamental nutritional niches (FNN) for maximal crop yield (Figure 1E) (34, 39).

**Figure 1:**
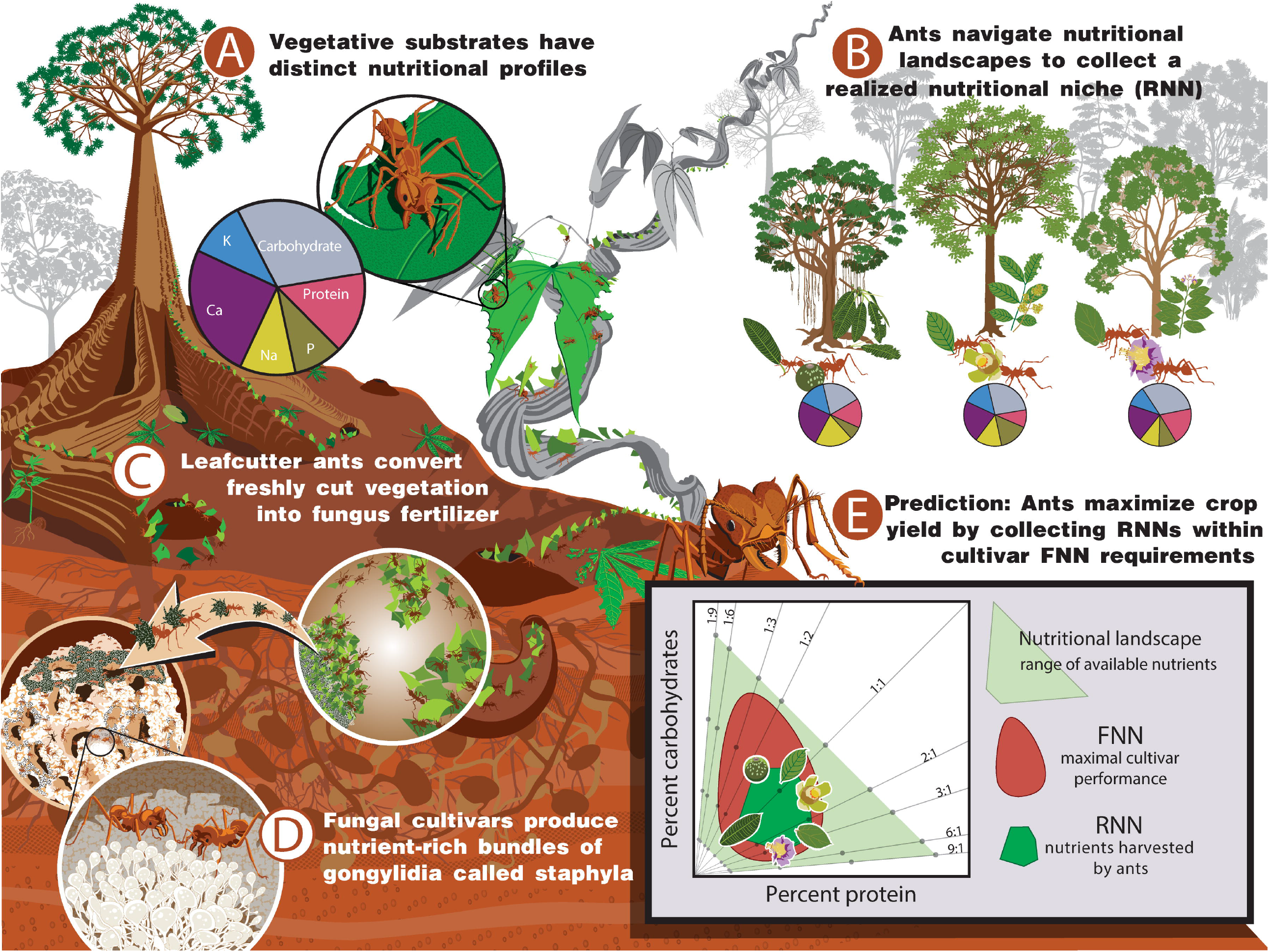
A niche-based framework for testing the hypothesis that leafcutter ants navigate tropical forests to collect plant substrates that target their fungal cultivar’s nutritional needs. (A) (A) Foragers can select among plant substrates (e.g. leaf, fruit, flower) that have distinct blends of protein, carbohydrates, and minerals. (B) Colonies can regulate nutritional intake by foraging across hundreds of plant species to acquire a realized nutritional niche (RNN). (C) Gardener ants convert foraged plant fragments into a nutritional mulch used to provision their fungal cultivar. (D) These nutrients promote hyphal growth and the production of edible nutrient-rich hyphal tips called gongylidia (packaged in bundles called staphylae). (E) We can study the ants’ nutrient provisioning strategy in two steps. We first define the cultivar’s fundamental nutritional niche (FNN) by measuring its performance when isolated onto petri dishes and grown across nutritional gradients, shown here as the light-green trapezoid ranging across protein:carbohydrate ratios (1:9 to 9:1 Pr:C) and protein + carbohydrate concentrations (4 diagonal rows of individual diet treatments (grey dots) with negative slopes ranging from 4 to 25 g/L). The red region indicates a hypothetical FNN of maximal cultivar performance. We then quantify the realized nutritional niche (RNN, dark green polygon) from nutrients contained in plant fragments foraged by free-ranging colonies. We array each plant fragment type based on their percent protein and carbohydrates and test the prediction that ants maximize cultivar performance by providing an RNN whose dimensions overlap with the cultivar’s FNN. The illustrations are by Damond Kyllo.

Recent lab-based experiments with nutritionally-defined diets have shown that: 1) *A. colombica* colonies tightly regulate protein foraging at low levels while allowing carbohydrate intake to fluctuate, and 2) the fungal cultivar is more sensitive to fluctuations in protein than carbohydrates, with reduced growth and survival when protein concentrations in available substrates exceed ca. 20% total dry mass (34). However, studies of free-ranging leafcutters have shown that some colonies preferentially forage N-rich leaves and thus likely target proteins built from N-rich amino acids (22, 40-42). This mixed evidence of protein regulation is likely due to the chemical complexity of field-collected vegetation relative to the controlled protein:carbohydrate diets used to assess the cultivar’s nutritional needs in the lab. Specifically, the minerals that likely vary across vegetation fragments, but which remain at low levels in lab diets, can influence the metabolic activity of fungi and their ability to access other nutrients in foods (43-47).

In the present study, we sought to explain how *A. colombica* leafcutter ants navigate a lowland Panamanian rainforest landscape of taxonomically and chemically diverse plant substrates, and whether the multidimensional foraging strategies of ant workers are mediated by the FNNs of their fungal cultivar *L. gongylophorus*. To do this, we first determined the cultivar’s FNN dimensions across interacting gradients of two macronutrients (protein, carbohydrates) and 10 minerals (Al, Ca, Cu, Fe, K, Mg, Mn, Na, P and Zn). We next quantified RNNs by identifying and nutritionally analyzing the vegetation fragments sampled from the mandibles of laden *A. colombica* foragers in the field. By overlaying RNNs atop of cultivar’s FNNs, we sought to determine the decisive nutrients and minerals regulated by leafcutter ants when provisioning their cultivars.

## Results

### Minerals shape the cultivar’s macronutrient requirements

To assess the effects of minerals on the cultivar performance, we first established a performance baseline by quantifying the cultivar’s FNNs for hyphal growth and staphyla density across an *in vitro* gradient of protein and carbohydrate (Pr:C) availability. This echoed recent findings (34), but also included lower nutritional concentrations to visualize the cultivar’s FNN across a broader range of plant substrates. Maximal hyphal growth occurred across a broad carbohydrate gradient up to 60% of macronutrient dry mass and with carbohydrate-biased Pr:C ratios ranging from 1:9 to 1:1 Pr:C (i.e. the red area in Figure 2A, Figure S1A, Tables S1-S2). Staphyla density was maximized across a narrower range of carbohydrates (up to 40%) but a wider range of protein (up to 30%). Staphyla density also had two distinct FNN peaks, with one in a carbohydrate-biased region below 1:3 Pr:C and another in a protein-biased region below 6:1 Pr:C (Figure 2B, Figure S1B-C, Tables S1-S2). These results indicate that both fungal traits are more sensitive to fluctuations in protein than carbohydrates, and that colonies have opportunities to use targeted doses of protein to selectively promote staphyla production.

**Figure 2:**
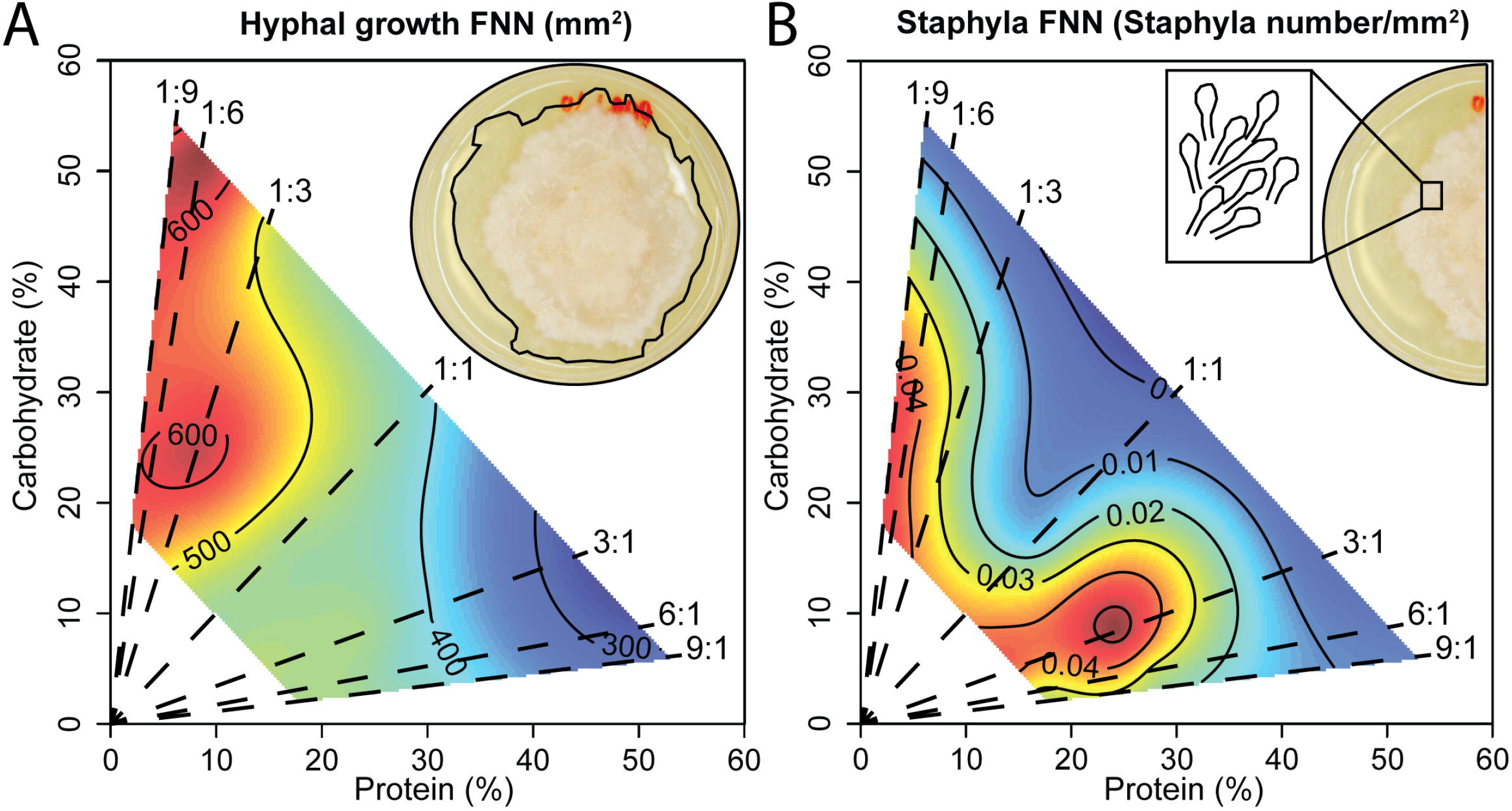
Quantifying the macronutrient FNN of the *L. gongylophorus* fungus cultivated by *A. colombica* leafcutter ants. A) Hyphal growth and B) staphyla density could both be maximized when provided carbohydrate-biased media, and both traits declined when protein-biased provisioning exceeded 30%. Staphyla density exhibited a second FNN peak at elevated protein concentrations (up to 30%) and relatively lower carbohydrate concentrations (up to 20%). Nutritional landscapes were generated by isolating *L. gongylophorus* from an *A. colombica* colony and performing *in vitro* experiments with nutritionally-defined media varying in protein:carbohydrate ratios (from 1:9 to 9:1 Pr:C) and protein + carbohydrate concentrations (4, 8 and 25 g/L).

We next examined the effects of minerals on cultivar growth relative to the baseline effects of macronutrients described above. We selected mineral addition treatments following extensive pilot experiments (Figure S2) and focused on hyphal growth as staphylae were often absent from mineral addition plates. Three minerals (Al, Fe, P) increased cultivar growth in the previously toxic protein-rich media, and P in particular was associated with up to 150% higher growth rates on the most protein-biased diets (Figure 3, Tables S3-S5). Other minerals either caused general toxicity effects by narrowing the cultivar’s FNN dimensions (Mn, Cu, K) or reducing cultivar growth across all protein and carbohydrate combinations (Ca, Mg, Na, and Zn) (Figure S3, Tables S3-S5). Fluctuations in mineral concentrations can thus reduce the cultivar’s growth performance and potentially render plant fragments with seemingly optimal blends of macronutrients unsuitable for cultivar provisioning.

**Figure 3:**
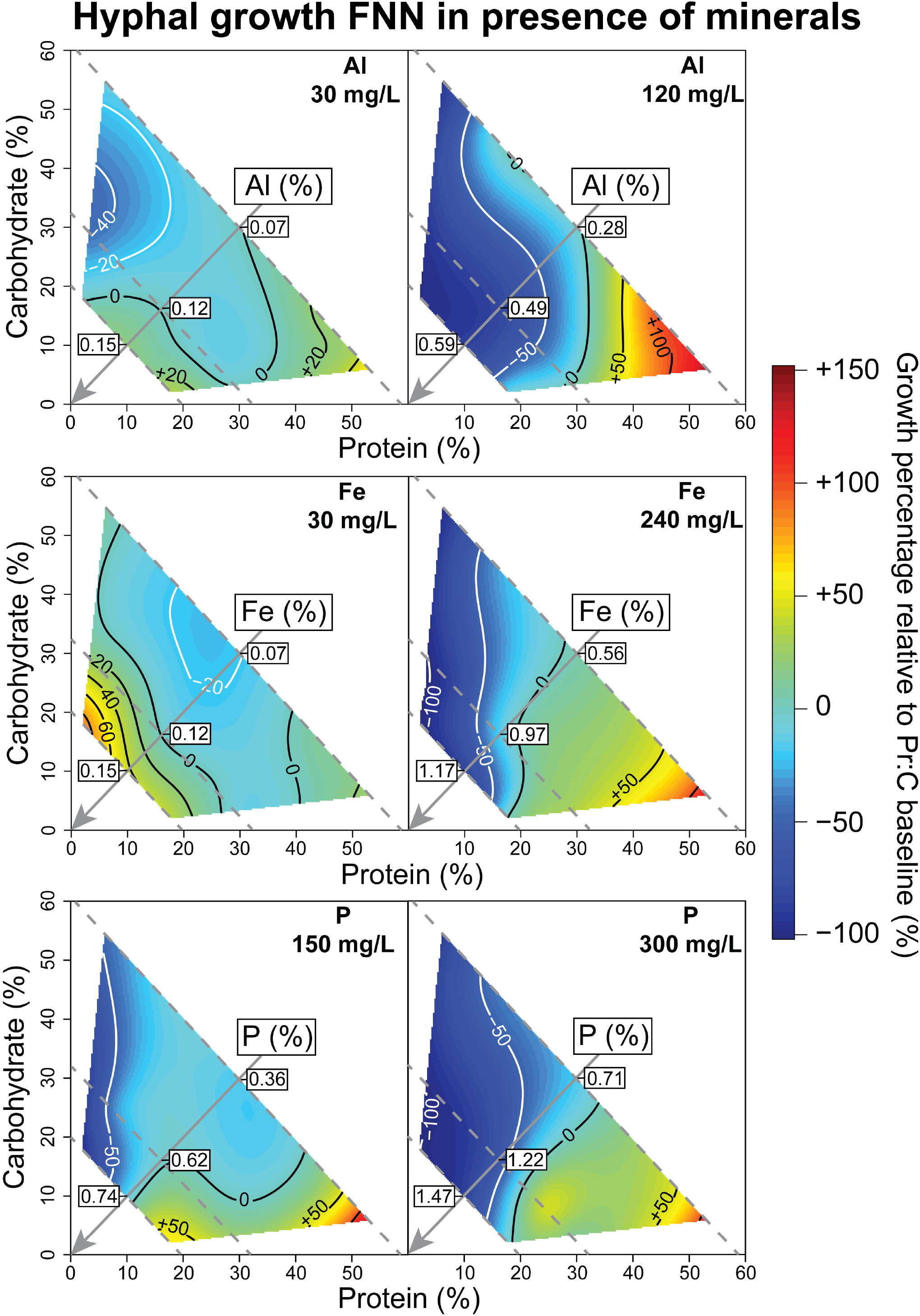
Quantifying the interacting effects of minerals and macronutrients on fungus cultivar growth. Three minerals (Al, Fe and P) expanded the FNN towards elevated growth in protein-rich conditions relative to baseline conditions without these minerals. Here we calculated relative growth percentage using the difference between cultivar’s final growth area in presence of each mineral relative to the same macronutrient condition without the mineral. The diagonal grey arrow indicates the gradient of mineral percent relative to protein and carbohydrate percent in diets. We provide two mineral concentration addition treatments. White isoclines indicate reduced growth relative to the macronutrient baseline, and black isoclines indicate increased growth. Seven other tested minerals (Ca, Cu, K, Mg, Mn, Na, Zn) induced varying degrees of toxicity for the cultivar across the gradient of protein and carbohydrate availability (see Figure S3).

### The macronutrient RNN targeted by free-ranging leafcutter ants

We next explored whether and how the cultivar’s FNN governs nutrient regulation strategies of foraging leafcutters in the field. We used Near Infrared Spectroscopy (48) to nutritionally analyze plant fragments carried by returning laden foragers from six *A. colombica* colonies (Table S6). We first quantified RNN dimensions in terms of protein and carbohydrates, based on collections of 44,533 plant fragments (dry mass 220.38 g) from 44 plant species (identified by DNA barcoding of ∼276 bp of *ITS1*; Figure 4A, Table S7) during 54 collection hours. Colonies foraged 2166 (± 283 SD) total fragments per 30-min, indicating that we collected ca. 40% (± 9% SD) of available fragments during each collection period (Figure S4). Colonies exploited similar numbers of plant species (Figure S5), although most plant species were foraged at low levels (Figure 4A, Figure S6, Table S8) and no plant species were common to all six colonies (Figure S5). Macronutrient concentrations in these substrates ranged from 5 to 42% carbohydrates and 5 to 35% protein (Figure 4B), yielding a broad RNN that overlapped with the cultivar’s FNNs for maximal hyphal growth (Figure 4C) and staphyla density (Figure 4D). Yet this RNN also exceeded protein levels that can reduce cultivar growth performance.

**Figure 4:**
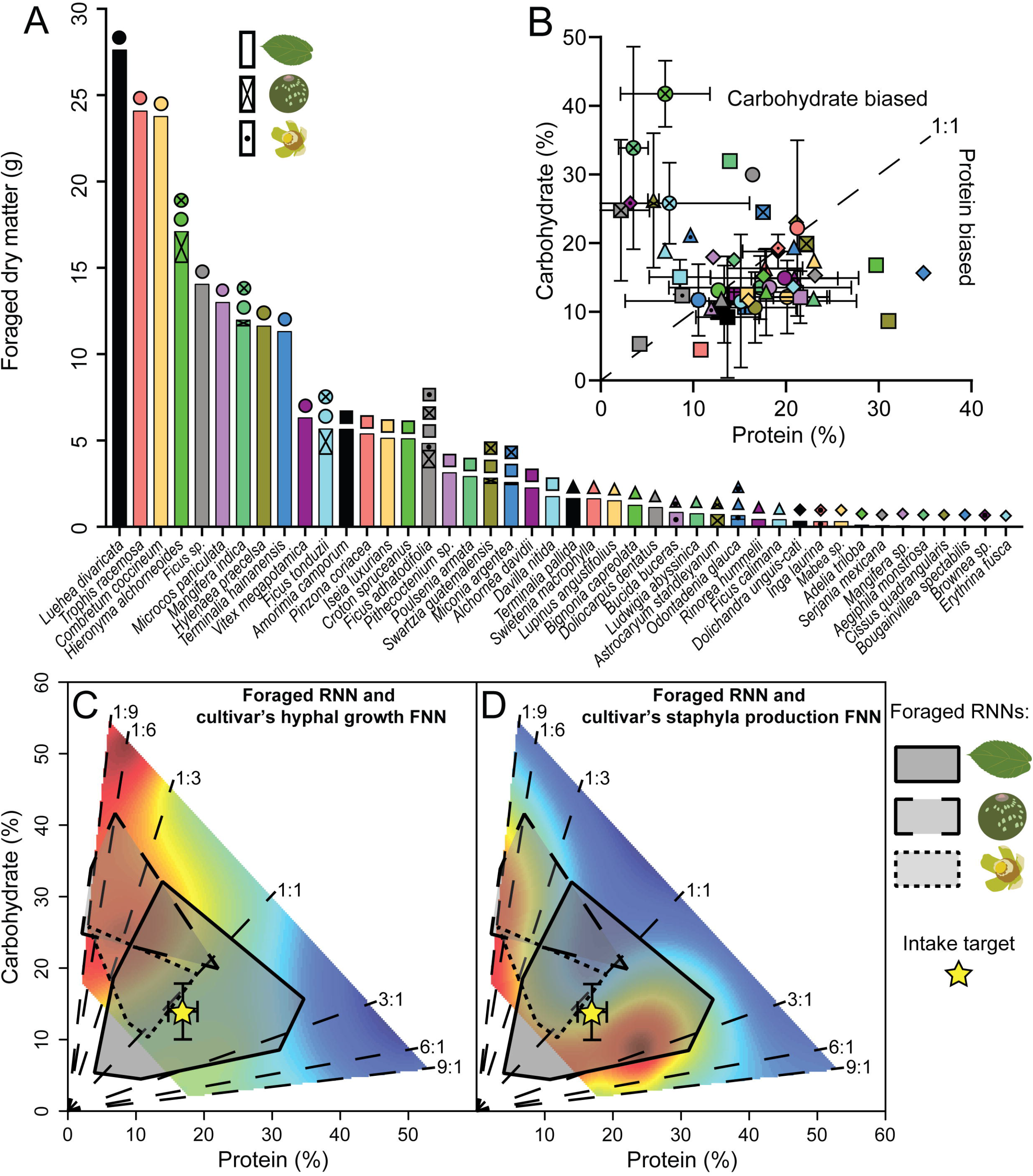
The plant substrates foraged by free-ranging leafcutter ants yield RNNs that overlap with their cultivar’s FNN requirements. (A) Colonies of *A. colombica* foraged mostly on leaves of a few plant species, but also collected leaves, fruits, and flowers from 44 plant species. (B) Foraged substrates (mean ± bidirectional SD) spanned macronutrient concentrations from 3 to 35% protein and from 5 to 42% carbohydrate. (C) The RNN polygon of leaves (n=40 species) contained protein in excess of the cultivar’s FNN for hyphal growth, and the RNNs of flowers (n=5 species) and fruits (n=7 species) both overlapped with the carbohydrate-rich FNN for maximal hyphal growth. (C) The leaf RNN overlapped with the protein-rich FNN for maximal staphyla density, and the fruit and flower RNNs overlapped with the carbohydrate-rich FNN for maximal staphyla density. The overall macronutrient intake target was slightly protein-biased (>1:1 Pr:C) and governed by leaf nutrients that contributed the majority of the foraged biomass. These results reflect the composite foraging of six *A. colombica* colonies observed during 54 collection hours. Results for individual colonies are provided in Figure S7-S8. Substrate illustrations in figure legends are by Damond Kyllo.

Colonies could target different RNN dimensions by collecting leaves (96.2% dry mass), flowers (2.9%), and fruit pieces (0.9%) (Figure 4A, Figure S6) as each substrate type had distinct blends of nutrients. First, while fruits and flowers had carbohydrate-biased RNNs, their RNNs did not overlap as fruits had higher carbohydrate concentrations (> 25%) than flowers (< 25%) (Figure 4B-C). Second, the leaf RNN spanned broader total macronutrient concentrations (5% to 50%) and tended to have more protein than flowers or fruits (Figure 4B-C). Leaves were also the dominant substrate type and thus governed each colony’s overall intake target, defined as the nutritional blend selected by a colony that in principle maximizes the cultivar’s performance, and against which surplus or deficient intake can be inferred (4, 5, 8). As a result, the intake target selected by ant foragers was biased towards protein levels that were beyond the cultivar’s FNN for maximal hyphal growth (Figure 4C, Figure S7), but near the protein-biased RNN for maximal staphyla density (Figure 4D, Figure S8).

### Optimal foraging requires multidimensional nutritional regulation

We next examined whether and how colonies regulate foraging of ten minerals to which the cultivar has narrow margins between limitation and toxicity. Mineral profiles varied widely across the 52 foraged plant substrates (Figure 5A, Table S8) and across leafcutter colonies (Figure S9, Table S9). Despite this variation, the more toxic minerals tended to occur at lower concentrations in foraged plant fragments (Figure 5B). Specifically, the most toxic minerals (Cu, Mn, Zn) that reduced cultivar growth at the lowest concentrations (60 mg/L) were measured in micrograms per gram of plant tissue (µg/g), while the least toxic minerals (Ca, K, Mg, Na) that the cultivar tolerated at higher concentrations (>600 mg/L) occurred in levels up to milligrams per gram (mg/g) of plant tissue (Figure 5B, Figure S2). The trace metals critical to an array of metabolic processes (Cu, Mn, Zn) thus appear to be generally foraged at lower levels than the ions that flux across cell membranes (Ca, K, Mg, Na). Despite these effects of individual minerals, the results thus far also show that mineral effects on cultivar performance are mediated by blends of macronutrients. We explore these nutritional interactions below.

**Figure 5:**
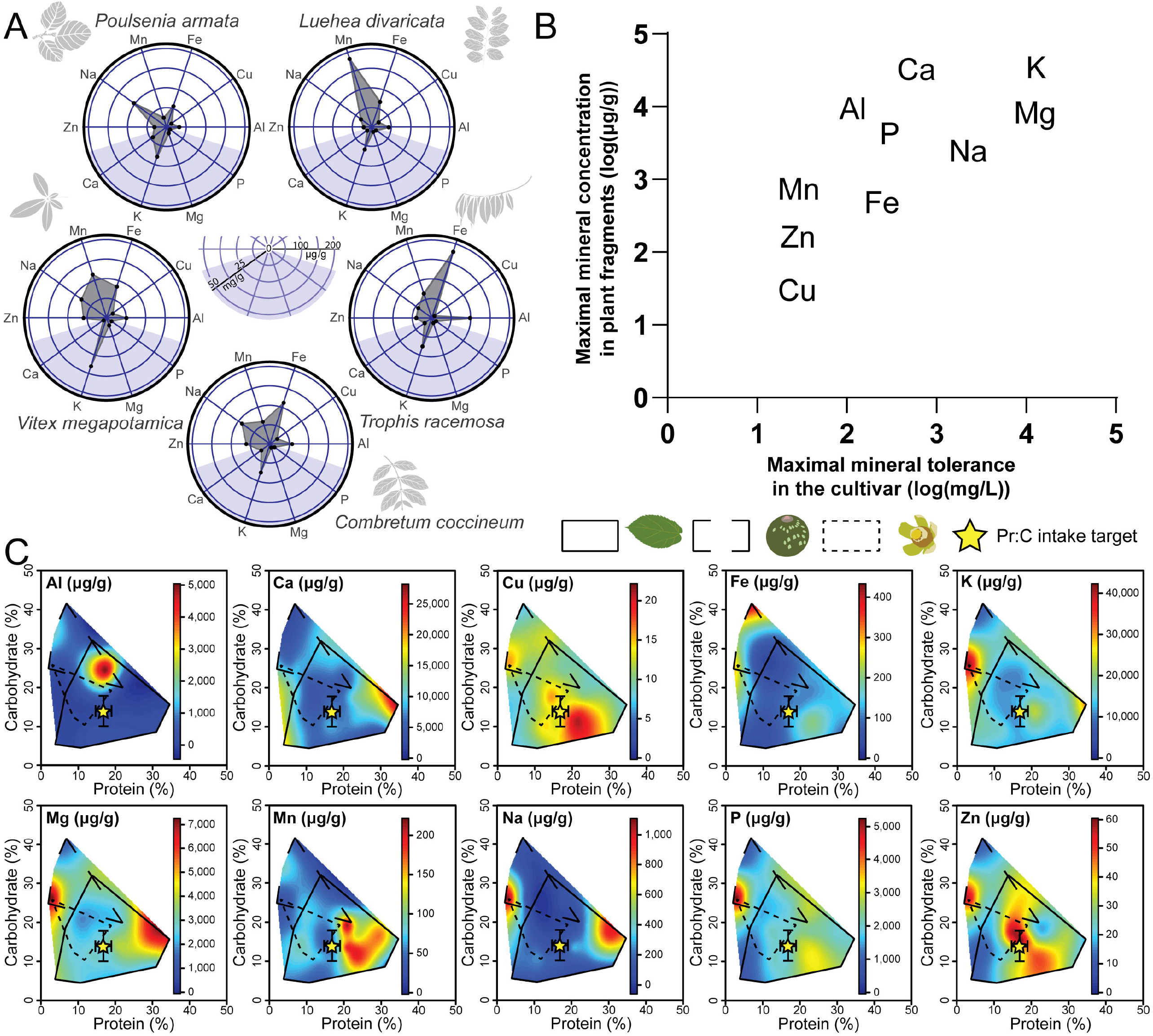
Testing for interactions between the mineral profiles and the macronutrient RNNs of foraged plant substrates. (A) Leaf mineral profiles of five foraged plant species illustrate the variation in concentrations of ten minerals observed across the 44 plant species. The minerals Ca, K, Mg, and P (blue shaded region of radial plots) are expressed in concentrations of mg/g, and Zn, Na, Mn, Fe, Cu, and Al are expressed in μg/g (white region). (B) There is a positive correlation between the cultivar’s maximal *in vitro* tolerance for each mineral and its maximal concentration in foraged plant fragments. (C) Mineral concentrations in foraged fragments are overlaid across gradients of protein and carbohydrates and interpreted relative to the RNN and intake target of these two macronutrients. See Results for at detailed interpretation. Substrate illustrations are by Damond Kyllo.

Solitary insect herbivores are known to tolerate more toxins when their diets also contain optimal blends of macronutrients that are close to the insects’ self-selected intake targets (49, 50). We extended this hypothesis to leafcutter ants and found that concentrations of the most toxic minerals (Cu, Mn, Zn) in foraged vegetation tended to peak closer to the macronutrient intake target compared to the less toxic minerals (Ca, K, Mg, Na) that peaked in carbohydrate-biased or protein-biased substrates (Figure 5C). This suggest that the negative effects of mineral surplus can be mitigated when the cultivar receives optimal blends of macronutrients. We next examined Al, Fe, and P that enhanced the cultivar’s protein tolerance, and hypothesized these minerals can expand a colony’s foraging niche across protein-biased plant substrates (Figure 3). We focused on the mineral RNNs for leaves, as fruits and flowers were never protein-biased and contributed little to total mineral provisioning due to their low biomass. As predicted, the most protein-rich leaves also contained the highest Fe and P concentrations (Figure 5C). The concentration of Al in leaves was not sufficient to assess this hypothesis as its RNN peak was due to a single fruit (Figure 5C). These results highlight that successful farming systems require multidimensional nutrient regulation and suggest a wealth of unexplored reciprocal adaptations in ants and fungal cultivars for optimizing this controlled provisioning.

## Discussion

The ecological success of leafcutter ants hinges upon nutritionally optimized fungus cultivar provisioning. We tested the hypothesis that ants regulate nutritional intake by foraging across plant species and substrate types to collect a realized nutritional niche (RNN) whose dimensions overlap with their cultivar’s fundamental nutritional niche (FNN) (Figure 1). We show that foraging must be optimized in N nutritional dimensions. First, a plant fragment can contain carbohydrate concentrations that maximize cultivar growth, and still be unsuitable for cultivar provisioning if its protein concentration exceeds ca. 20% (Figure 2). Second, this same protein-rich plant fragment may actually be suitable if it contains a narrow range of Al, Fe, or P that enhances the cultivar’s protein tolerance (Figure 3). We further show that colonies have opportunities to regulate their RNN dimensions by foraging across carbohydrate-biased fruits and flowers and protein-biased leaves (Figure 4). Yet, the cultivar’s narrow tolerance for fluctuating mineral provisioning also raises questions about the extreme generalist herbivory of leafcutter colonies that collect plant fragments with widely varying mineral profiles (Figure 5). More generally, the approach developed in this study provides a means to explain how the physiology of a microbial symbiont shapes to the multidimensional niche breadth of its insect host.

Like solitary generalist herbivores, leafcutter ants have the advantage of mixing many imbalanced foods to achieve balanced nutrition while also avoiding toxins (4, 51, 52). Unlike solitary herbivores, leafcutter colonies can recruit thousands of ants to sample many chemically diverse plant species at once (53). This distributed foraging facilitates dynamic nutritional regulation, but also poses unique challenges. Gardener ants in the nest must detect feedback from their fungal cultivar about its nutritional needs (54) and then communicate this information to foragers (55) and to other workers that dispose of large numbers of suboptimal fragments directly into enormous trash heaps (12, 56, 57). These sequential stages are analogous to pre-ingestive nutritional regulation of solitary herbivores (e.g. foragers select among imbalanced available plants) and post-ingestive regulation (e.g. gardeners expel nutritionally suboptimal fragments from the nest)(4). The critical regulatory decision points thus likely depend less on any decisive nutrient or nutrient-targeting enzyme (e.g. a pectinase) (58), but rather on the decisive cues (e.g. volatile semiochemicals (54, 59)) that enable ants to detect the cultivar’s immediate nutritional needs and then adjust provisioning.

Leafcutter ants also face different nutritional challenges than solitary herbivores. First, whereas most herbivores are protein limited (60), leafcutter ants tightly regulate protein intake if given the opportunity because excess protein can cause crop failure (34). Second, regulation of trace minerals is not typically assumed in insect herbivores (4), but fungal cultivars may experience minerals like plant quantitative plant defenses. Leafcutter colonies thus appear to forage the minerals with the highest potential for toxicity (Cu, Mn, Zn) at the lowest levels, and accept elevated concentrations only in leaves that are also optimized for macronutrient content. Minerals like Al, Fe, and P may further govern the leafcutter foraging niche, enhancing access to the most protein-biased leaves, perhaps by chemically immobilizing excess protein (61) or altering the cultivar’s metabolic processing capacity (46, 62, 63). In particular, the tight link between N and P content seen across plant species (64) suggests that the effects of P on the cultivar’s protein tolerance have been reinforced over millions of years of co-evolutionary crop domestication in *A. colombica* and other species of *Atta* that are also known to forage for N-rich leaves (*A. cephalotes* (22) and *A. laevigata* (41)).

Adaptative responses to mineral-macronutrient interactions would add to the impressive list of nutritional adaptations in the *L. gongylophorus* fungus, including expression of digestive enzymes that detoxify fresh vegetation (65) and degrade its carbohydrates (e.g. pectin) (66, 67). The cultivar’s physiological adaptations function seamlessly with the work of mobile gardener ants that ingest fungal gongylidia and then produce fecal droplets that distribute N-rich compounds (e.g. allantoin, ammonia, all 21 amino acids, (68)) and enzymes to begin digesting plant substrates on the growing fungus garden (38, 69). The nutritional contributions of bacteria within the farming symbiosis are also increasingly coming into focus, as they fix nitrogen (70), metabolize lipids (e.g. lipids, (71)), and can detoxify plant secondary metabolites (72). Within individual leafcutter ants, bacteria can also fix nitrogen (70) and metabolize citrate (33). Yet, leafcutter ants are the crown group of a monophyletic clade of over 250 fungus-farming ant species that scavenge mostly insect frass, tiny decaying wood pieces, flower bits, and occasionally mineral-rich insect cuticles for cultivar provisioning (32-34). The approach developed in the present study provides a means of linking the physiological traits of these diverse cultivars with the specific multidimensional nutritional challenges faced by foraging workers in complex foraging environments.

## Materials and Methods

### Fungal isolation and in vitro growth experiment

We isolated *L. gongylophorus* fungus gardens of two lab *A. colombica* colonies (ID: AC-2012-1 and AC-2014-2) collected in Soberanía Park (Panama) and maintained at the University of Copenhagen, in Denmark. We isolated fungus from the middle layer of the gardens and viewed the samples under a dissecting microscope. We then used a sterile dissecting needle to transfer staphyla to 60-mm petri dishes containing autoclaved potato dextrose agar media (PDA). Petri dishes were sealed with parafilm and incubated at 23.5 °C for a week in the dark. Clean fungal cultures were then transferred to new Petri dishes with PDA, sealed and incubated again for two weeks. We then isolated from these plates, repeating the transfer procedure, and letting these cultures grow for three weeks.

We used these isolates to estimate the growth rate of *L. gongylophorus* in a no-choice experiment with seven protein:carbohydrate diets (9:1, 6:1, 3:1, 1:1, 1:3, 1:6, 1:9 Pr:C) arrayed across three protein + carbohydrates concentrations (4, 8 or 25 g/L) (Table S10; adapted from (39). We added protein using equal amounts of bactopeptone (BD), bactotryptone (BD), and trypticase peptone (BD), carbohydrates using equal amounts of sucrose (Mamone) and starch (Sigma-Aldrich), bacteriological agar (VWR), and double-distilled water. Media were autoclaved at 121 °C and then 10 mL of were plated per sterile 60-mm petri dish under laminar flow. Plates were then exposed to UV light for 30 minutes. Fungus from PDA cultures was aseptically inoculated on each plate (n = 5 plates / diet) using a flame sterilized 4-mm diameter steel cylinder. We then sealed and stored all plates at 23.5 °C in the dark for 56 days during which we regularly checked plates and excised any contaminations. If plates were assessed to be heavily contaminated, we removed them from the experiment and inoculated new replicates. We outlined the outer edge of fungal expansion on each plate weekly.

We next assessed how ten minerals impact fungal performance, by adding the following compounds to the previously described media in solution: Ca (calcium chloride (CaCl_2_), Sigma-Aldrich), Na (sodium chloride (NaCl), Merck), K (potassium chloride (KCl), Sigma), Mg (magnesium sulfate (MgSO_4_), VWR), Zn (zinc sulfate heptahydrate (ZnSO_4_·7H_2_O), Sigma-Aldrich), Fe (iron sulfate heptahydrate (FeSO_4_·7H_2_O), Sigma), Cu (copper sulfate pentahydrate (CuSO_4_·5H_2_O), Sigma), Al (aluminum sulfate hydrate (Al_2_(SO_4_)_3_·H_2_O), Alfa Aesar), Mn (manganese chloride tetrahydrate (MnCl_2_·4H_2_O), Sigma-Aldrich), and P (85% phosphoric acid (85% H_3_PO_4_), Alfa Aesar) (Table S10). For some media (Al, Cu, Fe, P and Zn), the addition of minerals before sterilization resulted in permanent liquid diets after autoclaving. We thus added minerals to these diets after they were autoclaved to ensure solidification.

We initially performed a pilot study to identify experimentally relevant concentration ranges for each mineral. To do this, we inoculated and incubated plates as described above and observed them over 70 days. We used diet treatments including all seven Pr:C ratios at 8 g/L Pr+C concentration. We used eight concentrations for each mineral (n = 3 replicates per condition + 3 baseline plates with no mineral per condition; n = 1,890 plates). We chose three representative concentrations for each mineral: baseline (no mineral added), highest growth, and the highest concentration enabling growth. We then expanded the experiment to macronutrient concentrations of 4 and 25 g/L for the seven Pr:C ratios for the three concentrations for each mineral (Figure S10), replicating each mineral concentration 3 times with 3 baseline plates with no mineral (n = 1,260 plates).

### Measuring fungal performance

After the defined period of growth, we photographed each plate using a Canon EOS 7D Mark II camera mounted on a fix stand. We used ImageJ (v1.52a; (73)) to estimate fungal expansion (area, mm^2^) based on the final circumference line drawn around the outer border of the fungus using threshold contrast-adjusted greyscale images (with pixel_2_ = 0.02). We counted staphylae directly from plates viewed under a dissecting microscope for the no-choice diet experiment plates with no minerals. We used pheatmap package (v1.02.12; (74)) in RStudio v3.6.2 (75) to plot hyphal growth across the seven Pr:C ratios and 16 mineral concentrations for the 8 g/L Pr+C dilution. We used fields package (v10.3; (76)) in RStudio to plot cultivar FNNs for hyphal growth and staphyla density across nutritional landscapes with topological resolution of response surface, using λ = 0.001 as the smoothing parameter. To facilitate comparison with nutritional data attained from field-collected substrates (described below), nutritional concentrations in growth media were converted from g/L to % of total protein and carbohydrate mass relative to the total dry biomass of the growth media including non-nutritive components like agar.

### Substrate collections from A. colombica colonies

We located six colonies of *A. colombica* 287 m to 13.9 km apart in lowland tropical rainforest habitats at Soberanía National Park, Panama during the wet season (a period of high ant activity) from May 2 to June 29 2019 (Table S6). Vouchers of *A. colombica* ants were deposited in the Museo de Invertebrados Fairchild, Universidad de Panama. We then laid on trash bags next to each colony’s main nest entrance next to the most active foraging trail and collected plant substrates from laden returning foragers. Each collection event lasted 1.5 hours (between 9:00 and 12:00 AM) and was repeated by two observers on three days per colony for a total of 9 collection hours per colony. Each collection event included three 30-minute sampling periods, after which all collected substrates were placed into Ziploc bags and stored in a cooler. Total foraging levels on trails were then estimated by counting laden returning foragers during three observation periods of 10 minutes (using a manual counter) between each 30-minute sampling periods. These data showed that our substrate sampling capacity ranged from 39 to 50% of each colony’s total foraging effort (Figure S4). Back at the lab, we used a dissecting stereoscope to separate and weigh substrates based on morphospecies identifications. Substrates were further separated per colony and collection day, and then freeze-dried with a SP Scientific BenchTop Pro with Omnitronics for 24 hours. Samples were then weighed for dry mass and stored at -20°C in Ziploc bags with silica gel.

### DNA extraction and identification of plant samples

Back in Copenhagen, we homogenized ground freeze-dried plant samples in 10% Chelex (Sigma) and extracted DNA following 30 minutes of incubation at 100°C. We amplified by PCR the *Internal Transcribed Spacer 1* (*ITS1*) genetic marker using primers containing both the generic M13 sequences (used for subsequent sequencing) and the specific *ITS1*-specific Trac01 sequences (<u>M13F</u>-Trac01F 5’ <u>TGTAAAACGACGGCCAGT</u>GATATCCRTTGCCGAGAGTC 3’ and <u>M13R</u>-Trac01R 5’ <u>CAGGAAACAGCTATGAC</u>GAAGGAGAAGTCGTAACAAGG 3’). Sequencing was performed by Eurofins Genomics. We then performed a Blast-n with the DNA sequences in the NCBI database and attributed species identification to the best hit (based on E-value and percent identity). When a given sequence obtained several equally possible results, we restricted our identification to the genus level. We identified 44 different species from the 87 samples initially categorized into morphospecies (Table S7).

### Protein and carbohydrate composition of plant samples

We placed freeze-dried plant substrates in centrifuge tubes, plunged them into liquid nitrogen, and homogenised the samples using a plastic pestle. We used near infrared reflectance spectroscopy (NIRS) to estimate the concentrations of total nitrogen, and total non-structural carbohydrates (water soluble carbohydrates + starch) from the 87 substrates. We acquired NIRS spectra for each sample using an Antaris II FT-NIR Analyzer (Thermo Scientific) from 4.000 to 10.000 cm^-1^ (2.500 to 1.000 nm) at a resolution of 16 cm^-1^ and 2x gain. We used the standard default instrument calibration as the reference measurement. Each spectrum acquisition was the mean of 32 monochromatic scans. We performed three replicate spectrum acquisitions with repacking and calculated a mean spectrum for each sample. We selected a representative subset of samples for wet chemical analyses using Principal Component Analysis on centered NIRS spectra of the 87 initially identified sample types after pre-processing using 1^st^ derivative model on SIMCA software (Umetrics). We selected the samples according to their position on PCA axes (farthest away from the center of the data and within the large cluster of scores; (77)), and depending on whether we had sufficient biomass to meet the requirements of the chemical analyses.

We used a CN analyser (Eurovector) coupled to an isotope ratio mass spectrometer (Isoprime) to quantify total N from 3 to 4 mg of ground samples. We then estimated the quantity of total protein by multiplying total N by 6.25, a standard conversion approach in the literature (78). We estimated total non-structural carbohydrates (hereafter carbohydrates) by quantifying water-soluble carbohydrates with a Total Carbohydrate Assay Kit (Sigma-Aldrich) and starch with a Total Starch Assay Kit (Megazyme) using 25 and 50 mg of homogenized plant material, respectively. We used peach powder as a positive control and water as a negative control in these analyses. We used these wet chemical data to build partial least squares regression prediction models of the percentage of total protein and carbohydrates using the first derivative of the NIRS spectra in SIMCA software (Umetrics) (79). Prediction models were cross-validated with seven segments and permutation tests with 500 re-calculations were used to analyze models. Root mean square error of the estimation (RMSEE) for observations in the workset, root mean square error computed from the selected cross validation round (RMSECV), and R^2^ indicating the relationship between the measured and predicted samples were used to evaluate model performance ((80); Table S11).

We used the barcoding results to combine conspecific samples by calculating mean protein and carbohydrate values, and used these data to generate protein and carbohydrate RNNs for each of the six observed colonies (Figure S7-S8) and to generate a composite RNN for this population of *A. colombica*. We defined RNNs as the region bounded by each general plant substrate type (leaf, fruit, flower). The RNN from one of the colonies (colony 4) was previously published in (34) as part of a comparative analysis of fungus-farming ants and is included here in a different conceptual context and as part of a much-expanded dataset about the foraging ecology of *A. colombica*.

We calculated a macronutrient intake target by translating the substrate collection data into the actually foraged levels of protein and carbohydrates using arithmetic means weighted relative to total biomass (81). For each substrate we multiplied its percent protein or percent carbohydrates by the associated dry biomass (M). We then summed these values for each colony-observation period and divided this by the summed dry biomass of all substrates corresponding to colony-observation period. We used the following formula illustrated here for protein:

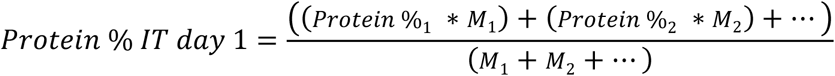

We then calculated colony-level intake targets by averaging intake targets across each colony’s three observation periods, and calculated a composite *A. colombica* intake target, by averaging across the six colony-level intake targets (Table S9).

### Mineral element composition of plant samples

We used a plastic pestle to homogenize freeze-dried plant substrate samples with available biomass ranging from 2 to 100 mg in centrifuge tubes after they were plunged into liquid nitrogen. We then weighed this plant material into Teflon microwave digestion tubes (n = 3 technical replicates per sample) and added 2.5 mL of 70% (v/v) nitric acid and 500 μL of 15% H_2_O_2_ for samples weighing > 20 mg, or 500 μL of 70% (v/v) nitric acid and 250 μL of 15% H_2_O_2_ for samples weighing < 20 mg. The tubes were then capped and the samples were digested in a microwave oven at 242 °C for 25 min (UltraWAVE single reaction chamber microwave digestion system, Milestone Inc.; CT Multiwave 3000, software version 1.24, Anton Paar GmbH). The resulting solutions were transferred to polypropylene vials and diluted with MilliQ water to a final volume of 50 mL for samples weighing > 20 mg or 10 mL for samples weighing < 20 mg. The elemental composition of these samples was measured for Al, Ca, Cu, Fe, K, Mg, Mn, Na, P and Zn using inductively coupled plasma optical emission spectrometry (ICP-OES; Agilent 5100, Agilent Technologies) (82, 83). Apple powder and a MilliQ water negative control were also analyzed as reference samples. For each substrate sample, we used technical replicates to calculate a mean value for each element. For each substrate type (leaf, fruit, flower) for each substrate species, we then calculated a mean (± SD) for each of the ten elements. Colony mineral intake targets were estimated by calculating the mean of the three weighted means (one for each day of collection) as described in previous paragraph (Figure S9, Table S9). We then mapped substrate mineral concentrations across gradients of substrate protein and carbohydrate concentrations using the fields package v10.3 (76) in RStudio with topological resolution of response surface λ = 0.001 as the smoothing parameter.

### Statistical analysis

Statistical analyses were performed in RStudio v1.2.5042 (75). We log-transformed the response variables when necessary to improve normality. We used least-square regressions to assess the underlying significance and interactions of both linear and quadratic terms and to support the interpretation of FNN heatmaps: 1) showing variation in fungus hyphal growth area and staphyla density across the 21 protein and carbohydrate diet combinations (Tables S1-S2), and 2) showing hyphal growth area across the 441 protein:carbohydrate:mineral substrate combinations with a separate model for each mineral (Tables S3-S5). Venn diagrams of foraged plant species distribution across habitats and colonies were generated using eulerr R package (84). The overall dataset and supporting R scripts are available as supplementary files.

## Supporting information

R_Script

## Acknowledgments

Thomas Hesselhøj Hansen and Lena Asta Byrgesen from the Department of Plant and Environmental Sciences of the University of Copenhagen performed the ICP-EOS analysis. We thank Damond Kyllo for figure illustrations, Audrey Dussutour for advice with nutritional analyses, and Jacobus Boomsma for comments on an earlier draft of the manuscript. We thank the Smithsonian Tropical Research Institute for logistical support during fieldwork. This research was funded by an European Research Council Starting Grant (ELEVATE: ERC-2017-STG-757810) to J.Z.S. The Ministerio de Ambiente, Republica de Panama provided permits for field research (SE/A-24-19) and sample exportation (SEX/A-41-19). The authors have declared no competing interest.

