## Supplementary information for "The multidimensional nutritional niche of fungus-cultivar provisioning in free-ranging colonies of a neotropical leafcutter ant"

**This PDF file includes:**

Figures S1 to S9
Tables S1 to S11
Legends for Dataset1

**Other supplementary materials for this manuscript include the following:**

Dataset1
R_Script

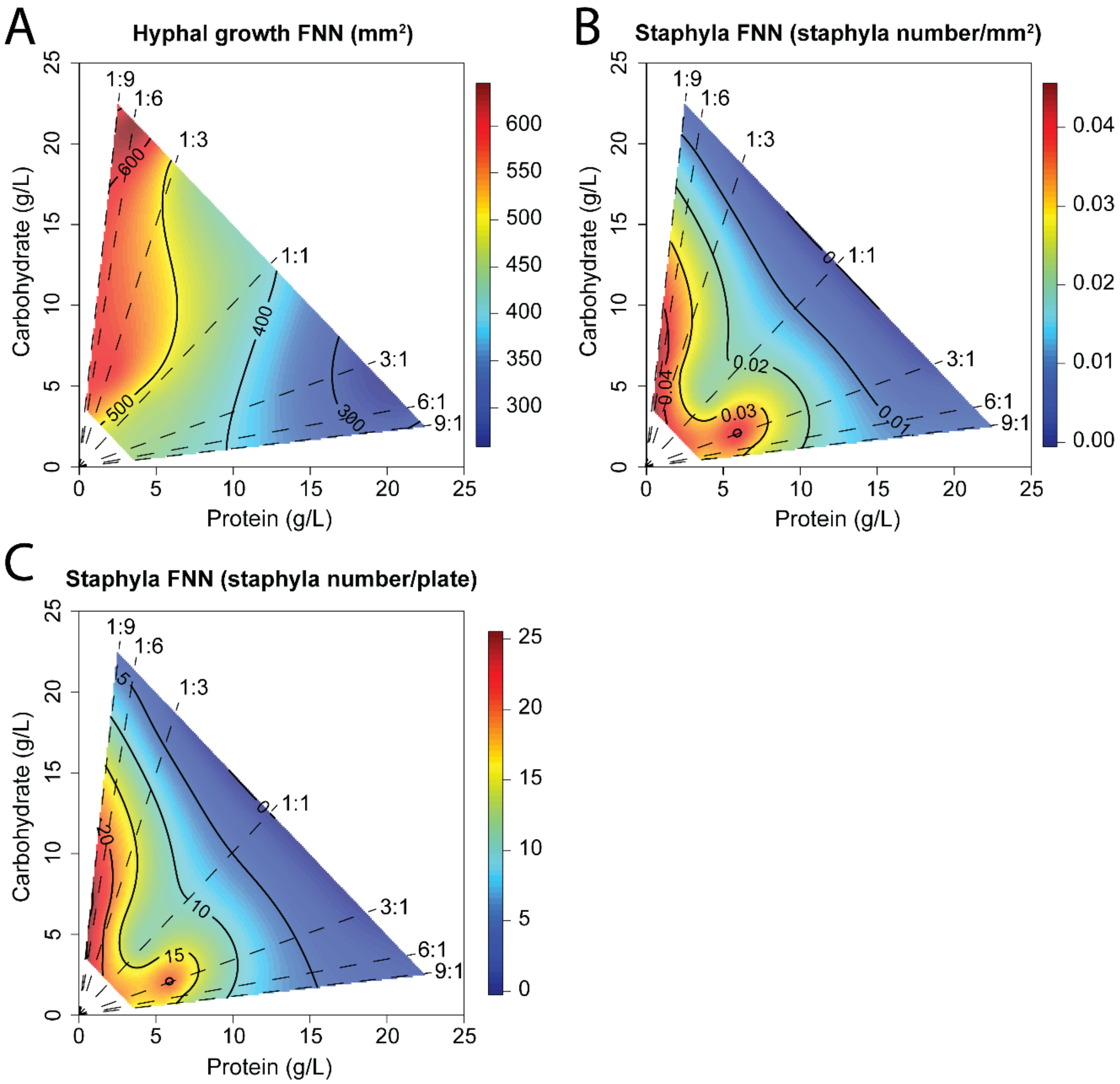


**Figure S1: Quantifying the fundamental nutritional niche (FNN) of the *Leucoagaricus gongylophorus* fungal cultivar in terms of protein and carbohydrate levels expressed in g/L**. These heatmaps reflect the results shown in Figure 2 but are expressed in terms of g/L macronutrient dilutions in the *in vitro* diet media rather than percent protein and carbohydrates relative to dry media mass. Percent macronutrient concentrations were used to compare fungal FNNs with RNNs provided by nutrients in plant substrates (Figure 4). (A) The hyphal growth of the fungal cultivar is maximized across a wider range of carbohydrates than protein (B, C) The density and number of staphylae per plate is maximized on carbohydrate-biased diet below 15 g/L carbohydrates and 3 g/L protein, and on protein-biased diets below 4 g/L carbohydrates and 8 g/L protein.

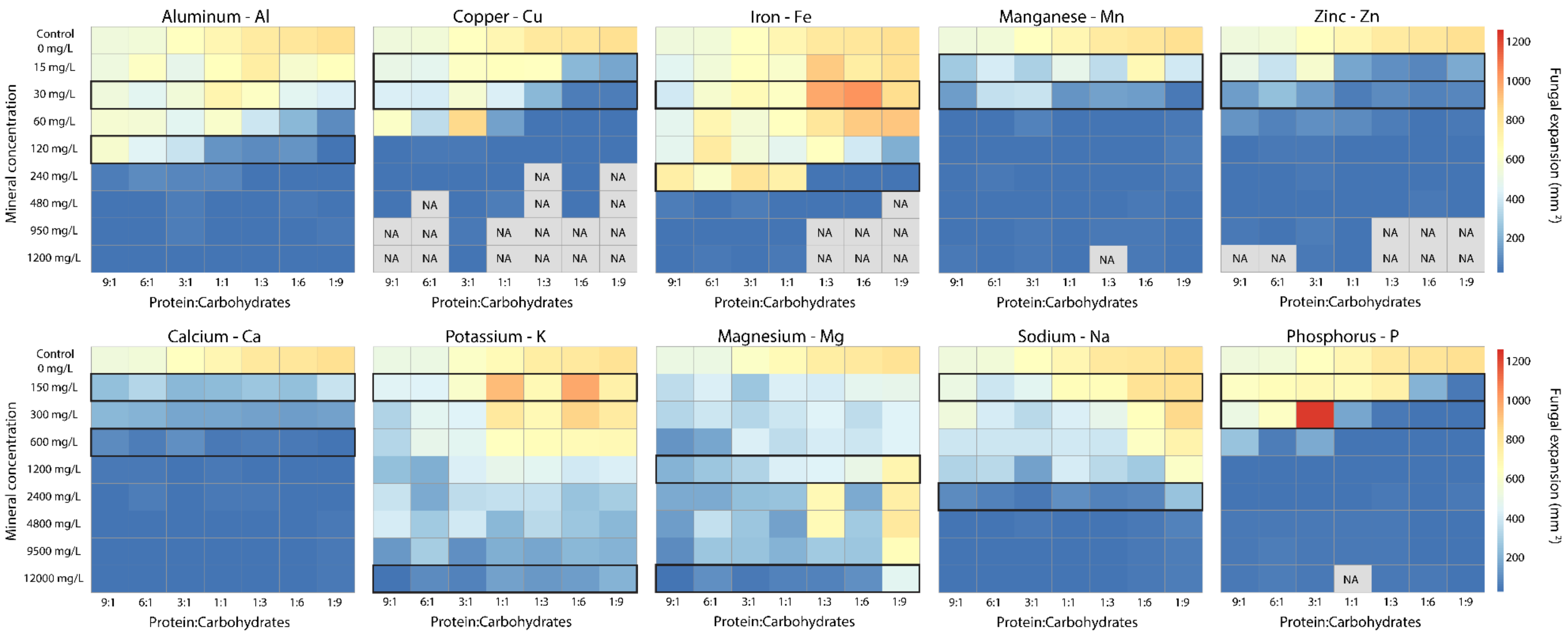


**Figure S2: Pilot studies showing hyphal growth in the *Leucoagaricus gongylophorus* fungal cultivar across 7 Pr:C ratios and 16 mineral element concentrations at the 8 g/L Pr+C dilution.** The cultivar exhibited lowest tolerance to minerals Cu, Mn, and Zn, with substantially reduced growth when these minerals exceeded 60 mg/L. The cultivar was able to sustain growth at slightly higher concentrations for Al (120 g/L) and Ca (600 g/L). The cultivar exhibited somewhat higher tolerance as the other minerals increased in concentration, and even had increased growth relative to the macronutrient baseline with Fe, Al, and P on protein-biased diets. For each mineral, two mineral-concentration rows have a black outline indicating they were used in the scaled-up experiment presented in this manuscript at 4 and 25 g/L Pr+C dilutions. These two rows reflect: 1) the mineral concentration enabling the highest cultivar growth, and 2) the highest mineral concentration enabling cultivar growth. NA indicates that data were not-available because media did not solidify after autoclaving (see methods) making culture impossible. Since we observed toxicity at lower mineral concentrations, we did not repeat these high-concentration treatments.

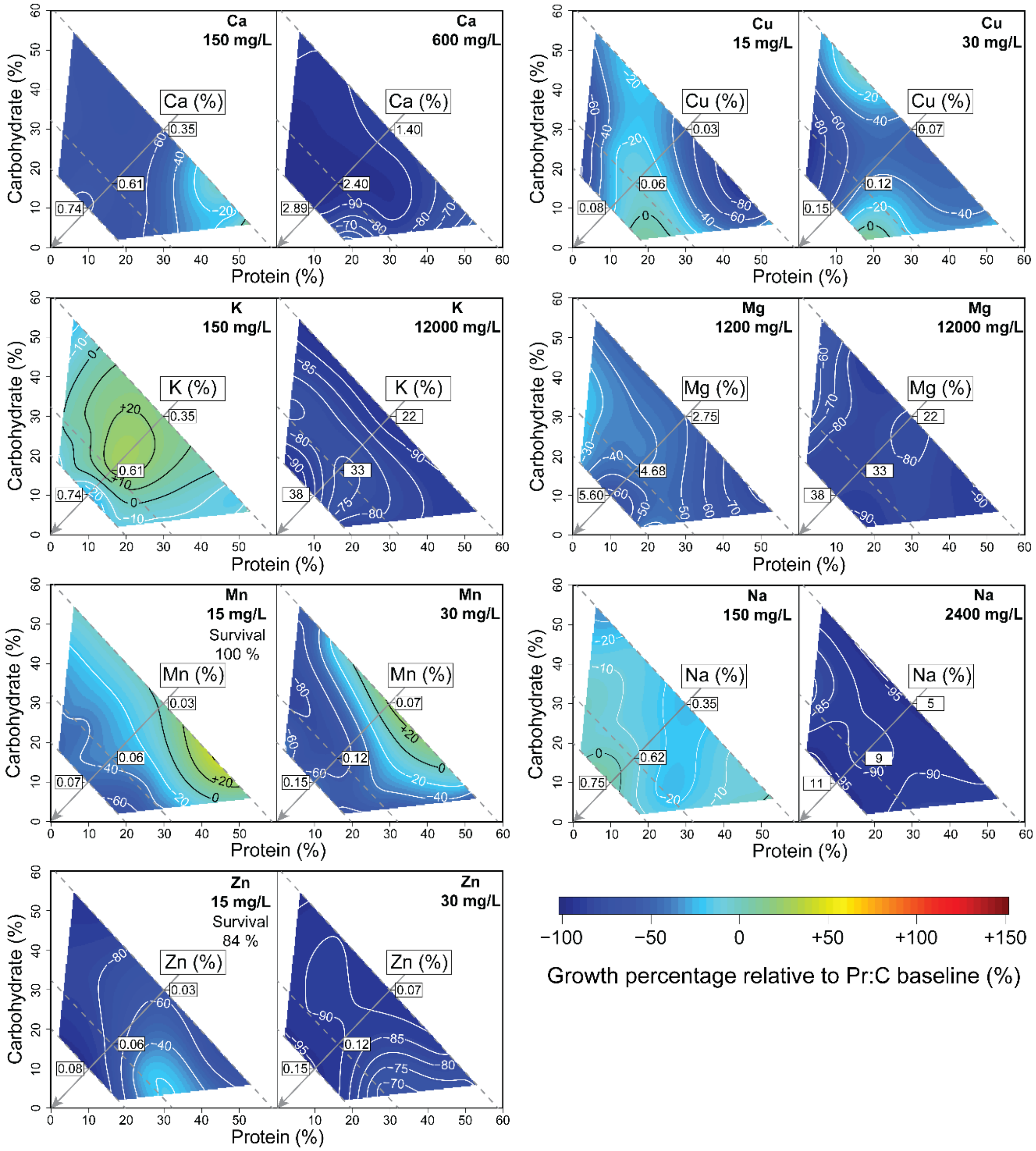


**Figure S3: Quantifying the interacting effects of minerals and macronutrients on fungus cultivar growth.** We calculated the relative growth percentage using the difference between the area of fungus growth in presence of mineral and its growth at the same Pr:C and Pr+C condition with no mineral. The diagonal grey arrow indicates the gradient of percentage of mineral relative to the percentage of protein + carbohydrates in diets. Some minerals (Cu, K, Mn) narrowed the cultivar's macronutrient FNN, while other minerals (Ca, Na, Mg, Zn) were toxic to the cultivar across all the protein and carbohydrate combinations. White isoclines indicate negative growth relative to the macronutrient baseline, and black isoclines indicate positive growth.

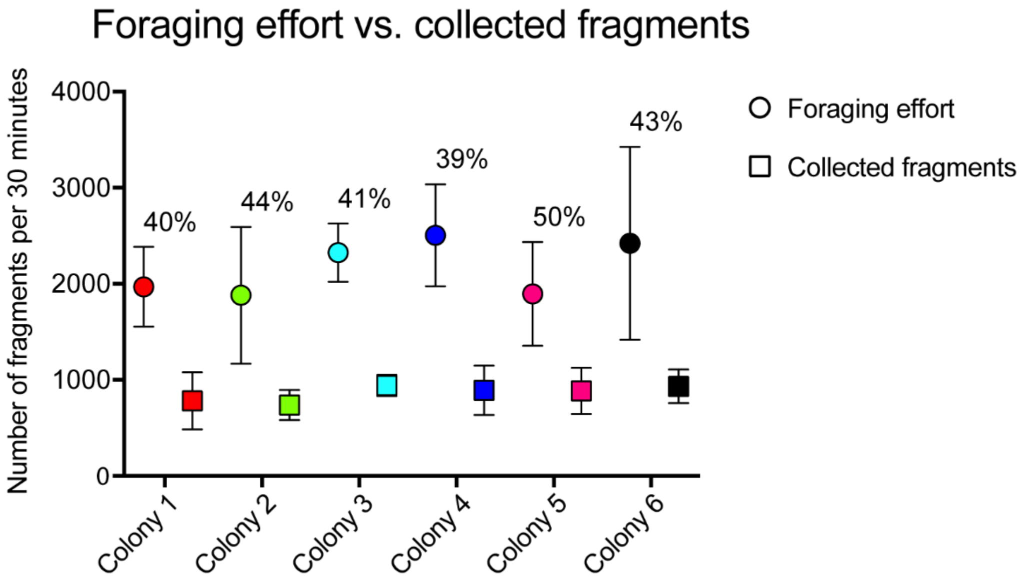


**Figure S4: Assessing the extent to which our plant fragment collections represented a colony's total foraging effort.** The variable 'foraging effort' is the mean (± SD) number of total plant fragments carried by returning laden workers during 10-minute observation periods and multiplied by 3 to estimate total activity during the 30-minute intervals in which we collected substrates. The variable 'collected fragments' is the mean number of plant fragments we actually collected from the mandibles of laden foragers from each colony during 30-minute sampling periods. Colony activity ranged between ca. 2000 and 3000 foraged items per colony, and sampling capacity was typically ca. 1000 items. The percentage provided for each colony indicates the proportion of fragments that were collected. Observation periods occurred on 3 separate days for each colony.

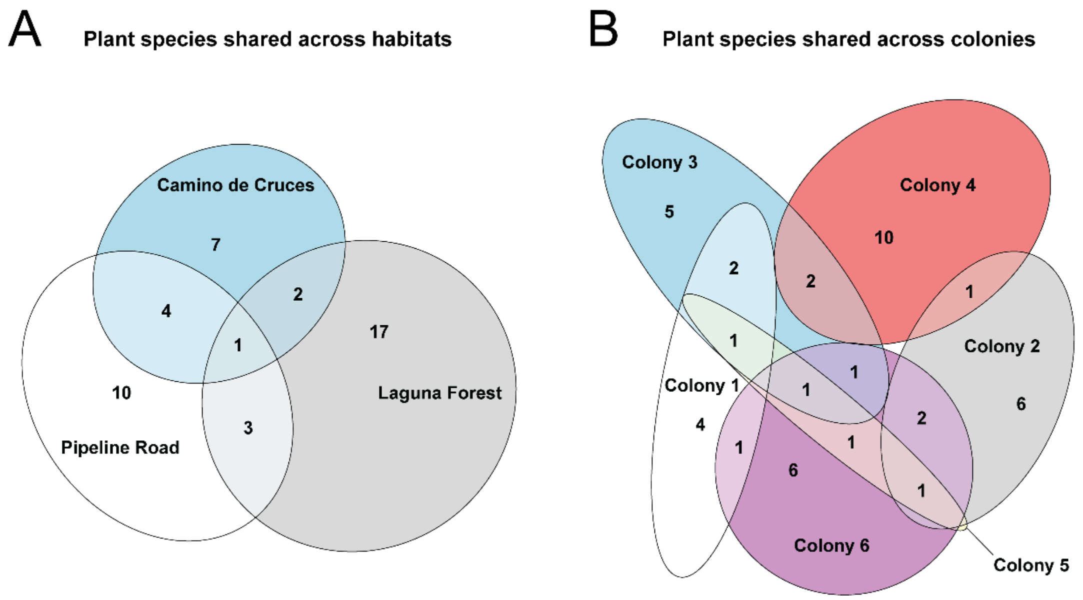


**Figure S5: Colonies collected 52 different substrates from 44 plant species, with many substrates unique to (A) habitats and (B) colonies.** Colonies 1 (8 species) and 2 (10 species) were located on Pipeline Road, colonies 3 (12 species) and 4 (13 species) were located in Laguna Forest and colonies 5 (4 species) and 6 (13 species) were from a forest called Camino de cruces. GPS coordinates are provided in Table S7.

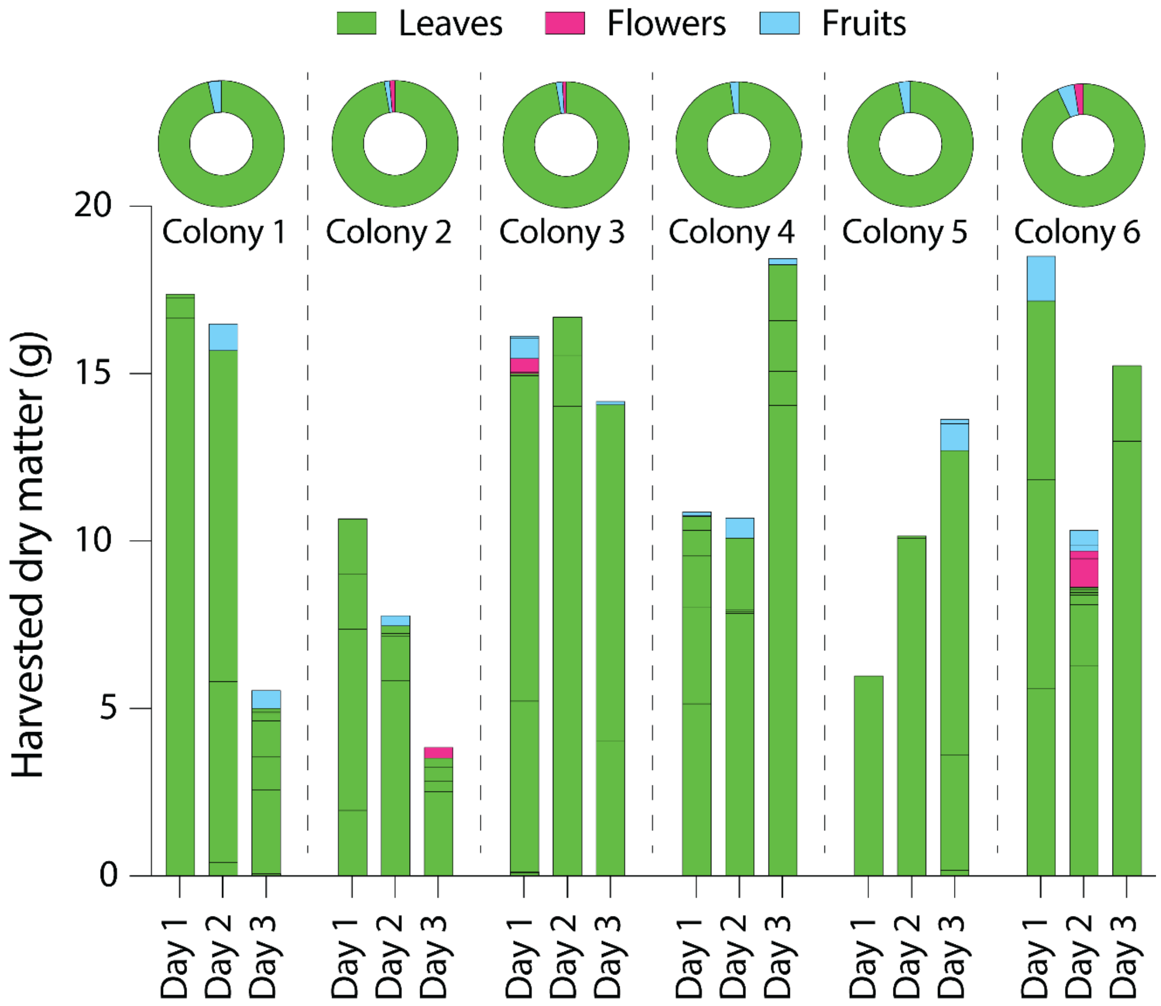


**Figure S6: Quantities and types of plant material collected by six colonies during three sampling days per colony and 54 total observation hours.** Colonies of *A. colombica* in this study collected mostly leaves with a small proportion of flowers and fruits. Individual colonies exhibited substantial variation in sampling effort across collection days. Each individual bar is a stacked bar showing the quantities of different plant species and substrates types. The circular pies represent the proportion of the three different substrates (leaves, flowers and fruits) collected per each colony across the three days of collection.

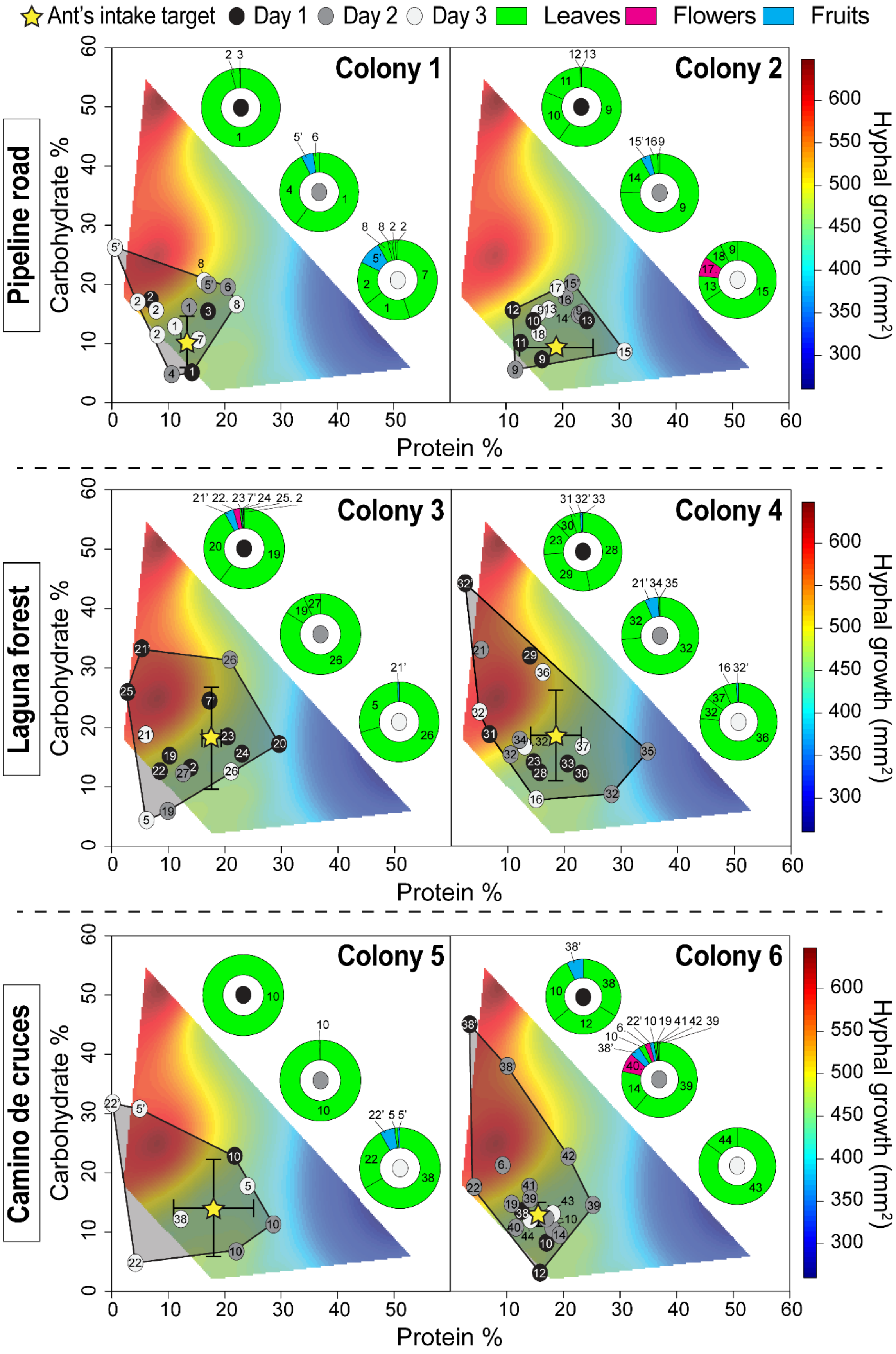

Ant's intake target
Day 1
Day 2
Day 3
Leaves
Flowers
Fruits
Pipeline road
Laguna forest
Camino de cruces
Colony 1
Colony 2
Colony 3
Colony 4
Colony 5
Colony 6
Carbohydrate %
Protein %
Hyphal growth (mm²)

**Figure S7: Comparing colony-level RNNs to the hyphal growth FNN requirements of the *L. gongylophorus* cultivar.** Pie graphs show the proportion of each foraged substrate type (leaf, flower, fruit) and species (1-44) per colony (1-6), per foraging day (1, 2, 3). Symbols used to construct each RNN contain numbers reflecting each plant species (1-44), with some numbers replicated within a given RNN polygon across multiple nutritional measurements (see methods), and with substrate types indicated as follows: 1 leaf, 1’ fruit, 1. flower. Numerical codes for species are as follows: 1 *Luehea divaricata*, 2 *Davilla nitida*, 3 *Adelia triloba*, 4 *Pinzona coriacea*, 5 *Ficus tonduzii*, 6 *Odontadenia glauca*, 7 *Miconia argentea*, 8 *Dolichandra unguis-cati*, 9 *Hylenaea praecelsa*, 10 *Combretum coccineum*, 11 *Terminalia pallida*, 12 *Amorimia camporum*, 13 *Rinorea hummelii*, 14 *Pithecoctenium sp.*, 15 *Swartzia guatemalensis*, 16 *Bignonia capreolata*, 17 *Inga laurina*, 18 *Mabea sp.*, 19 *Terminalia hainanensis*, 20 *Croton spruceanus*, 21 *Astrocaryum standleyanum*, 22 *Ficus adhatodifolia*, 23 *Swietenia macrophylla*, 24 *Serjania mexicana*, 25 *Brownea sp.*, 26 *Trophis racemose*, 27 *Doliocarpus dentatus*, 28 *Iseia luxurians*, 29 *Poulsenia armata*, 30 *Ludwigia abyssinica*, 31 *Ficus calimana*, 32 *Mangifera indica*, 33 *Erythrina fusca*, 34 *Mangifera sp.*, 35 *Bougainvillea spectabilis*, 36 *Ficus sp.*, 37 *Lupinus angustifolius*, 38 *Hieronyma alchorneoides*, 39 *Vitex megapotamica*, 40 *Bucida buceras*, 41 *Aegiphila monstrosa*, 42 *Cissus quadrangularis*, 43 *Microcos paniculate*, 44 *Alchornea davidii*.

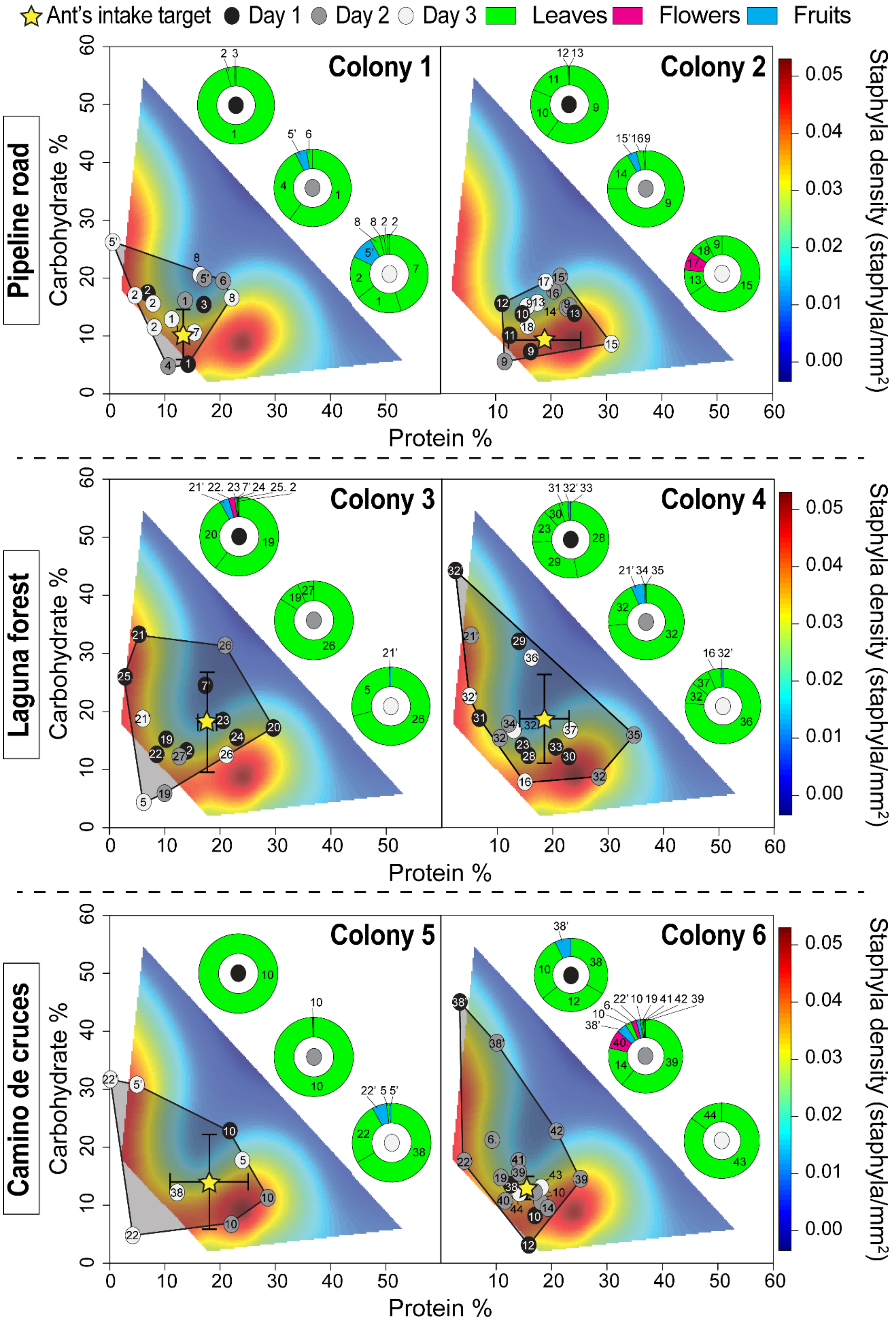
Ant's intake target
Day 1
Day 2
Day 3
Leaves
Flowers
Fruits
Pipeline road
Laguna forest
Camino de cruces
Colony 1
Colony 2
Colony 3
Colony 4
Colony 5
Colony 6
Carbohydrate %
Protein %
Staphyla density (staphyla/mm²)
0.05
0.04
0.03
0.02
0.01
0.00

**Figure S8: Comparing colony-level RNNs to the gongylidia density FNN requirements of the *L. gongylophorus* cultivar.** Pie graphs show the proportion of each foraged substrate type (leaf, flower, fruit) and species (1-44) per colony (1-6), per foraging day (1, 2, 3). Symbols used to construct each RNN contain numbers reflecting each plant species (1-44), with some numbers replicated within a given RNN polygon across multiple nutritional measurements (see methods), and with substrate types indicated as follows: 1 leaf, 1' fruit, 1. flower. Numerical codes for species are as follows: 1 *Luehea divaricata*, 2 *Davilla nitida*, 3 *Adelia triloba*, 4 *Pinzona coriacea*, 5 *Ficus tonduzii*, 6 *Odontadenia glauca*, 7 *Miconia argentea*, 8 *Dolichandra unguis-cati*, 9 *Hylenaea praecelsa*, 10 *Combretum coccineum*, 11 *Terminalia pallida*, 12 *Amorimia camporum*, 13 *Rinorea hummelii*, 14 *Pithecoctenium sp.*, 15 *Swartzia guatemalensis*, 16 *Bignonia capreolata*, 17 *Inga laurina*, 18 *Mabea sp.*, 19 *Terminalia hainanensis*, 20 *Croton spruceanus*, 21 *Astrocaryum standleyanum*, 22 *Ficus adhatodifolia*, 23 *Swietenia macrophylla*, 24 *Serjania mexicana*, 25 *Brownea sp.*, 26 *Trophis racemose*, 27 *Doliocarpus dentatus*, 28 *Iseia luxurians*, 29 *Poulsenia armata*, 30 *Ludwigia abyssinica*, 31 *Ficus calimana*, 32 *Mangifera indica*, 33 *Erythrina fusca*, 34 *Mangifera sp.*, 35 *Bougainvillea spectabilis*, 36 *Ficus sp.*, 37 *Lupinus angustifolius*, 38 *Hieronyma alchorneoides*, 39 *Vitex megapotamica*, 40 *Bucida buceras*, 41 *Aegiphila monstrosa*, 42 *Cissus quadrangularis*, 43 *Microcos paniculate*, 44 *Alchornea davidii*.

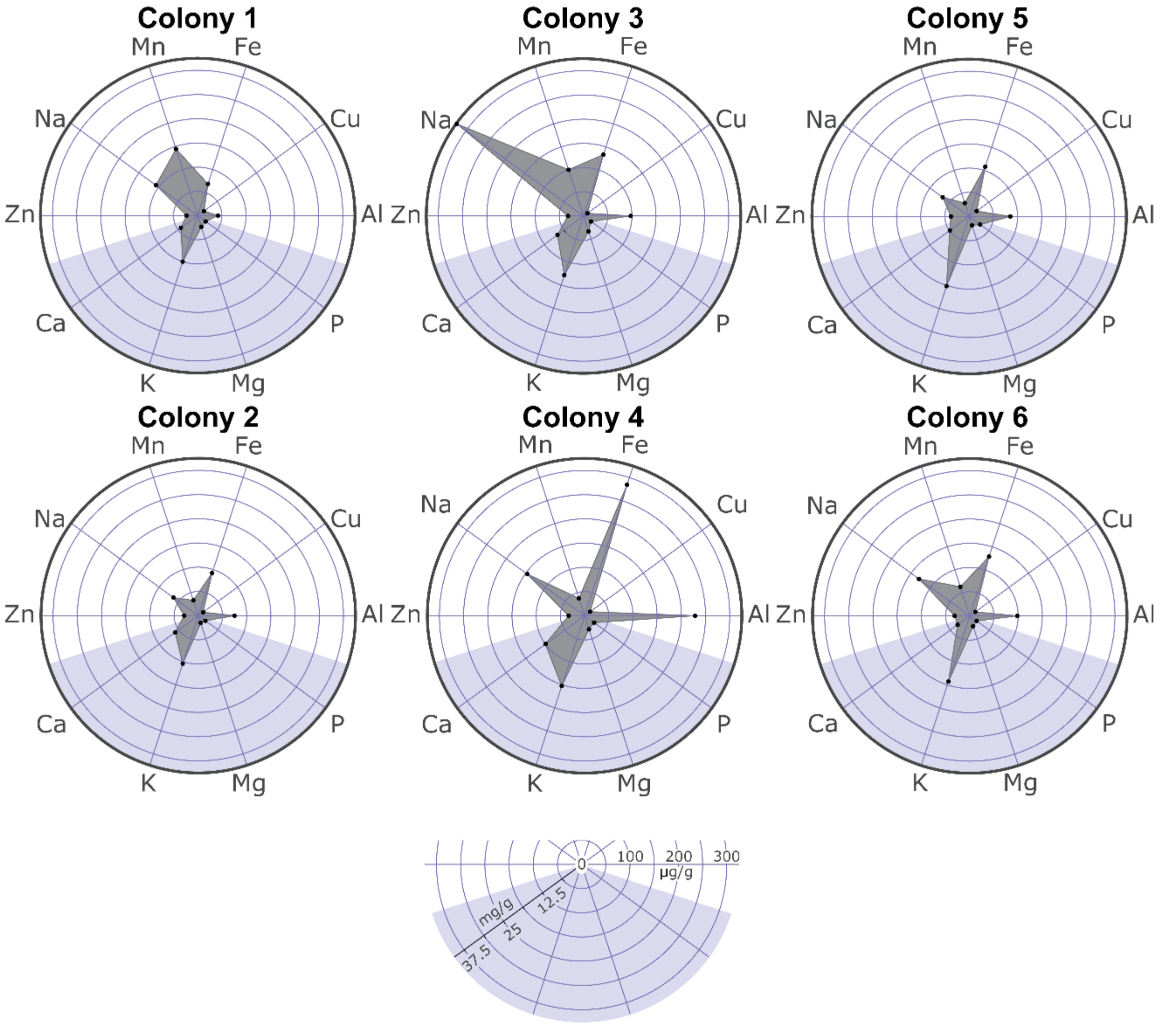


**Figure S9: Multidimensional mineral RNNs collected by each colony.** The intake target of each mineral is calculated using the weighted mean concentration for each of the three days of collection per colony to account for the difference in dry mass for each substrate collected. These mineral RNNs reflect a high degree of variation across colonies resulting from the complexity of plant material composition.

**Table S1: Univariate tests of parameter estimates and coefficients for hyphal growth and staphyla density across the protein-carbohydrate diet treatments in the *in vitro* experiment.**

| Response variable | Explanatory variables | Degrees of freedom | F (P) | β (SE) |
|---|---|---|---|---|
| Log(Hyphal growth) | Protein | 1 | 62.6727 (**3.62e-12**)*** | -0.0164414 (0.0333359) |
| | Carbohydrates | 1 | 4.7003 (**0.03255**)* | 0.0579577 (0.0333316) |
| | Protein$^2$ | 1 | 1.0403 (0.31025) | -0.0002956 (0.0011736) |
| | Carbohydrates$^2$ | 1 | 0.2934 (0.58927) | -0.0015489 (0.0011732) |
| | Protein x Carbohydrates | 1 | 1.8713 (0.17442) | -0.0031164 (0.0022781) |
| | Residuals | 99 | | |
| Staphyla density | Protein | 1 | 20.0118 (**2.057e-05**)*** | -2.458e-03 (2.672e-03) |
| | Carbohydrates | 1 | 19.7783 (**2.277e-05**)*** | 1.373e-03 (2.672e-03) |
| | Protein$^2$ | 1 | 2.7096 (0.1029) | 5.518e-05 (9.408e-05) |
| | Carbohydrates$^2$ | 1 | 0.7566 (0.3865) | -1.067e-04 (9.405e-05) |
| | Protein x Carbohydrates | 1 | 0.5330 (0.4671) | -1.333e-04 (1.826e-04) |
| | Residuals | 99 | | |

Diet treatments included 21 diets varying in the ratio (9:1, 6:1, 3:1, 1:1, 1:3, 1:6, 1:9 Pr:C) and concentration (4, 8, and 25 g/L) of protein and carbohydrates. The results shown here are from least-square regressions estimating relationships between diet treatments and hyphal growth and staphyla density. We include linear and quadratic coefficients for protein and carbohydrates in diets and their interactions. F indicates F-value and P the probability of F. P-values < 0.05 are in bold. β indicates coefficient of the model for each parameter and SE indicates standard error. Significance of estimated coefficients: *P < 0.05, **P < 0.01, ***P < 0.001.

**Table S2: Significance tests for univariate models for hyphal growth and staphyla density across Pr:C nutritional landscape.**

| | Model – Hyphal growth | Model – Staphyla density |
|---|---|---|
| **Degrees of freedom model** | 5 | 5 |
| **Degrees of freedom residuals** | 99 | 99 |
| **Adjusted $R^2$** | 0.3867 | 0.2717 |
| **Mean Square Model Residuals** | 0.0975 | 0.0006264 |
| **F-statistic** | 14.12 | 8.758 |
| **P-value** | 2.049e-10*** | 6.605e-07*** |

Diet treatments included 21 diets varying in the ratio (9:1, 6:1, 3:1, 1:1, 1:3, 1:6, 1:9 Pr:C) and concentration (4, 8, and 25 g/L) of protein and carbohydrates. Shown are the characteristics and significance of models from Table S1. Significance of estimated coefficients: *P < 0.05, **P < 0.01, ***P < 0.001.

**Table S3: Univariate tests of parameter estimates for hyphal growth in presence of mineral elements.**

| Response variable | Explanatory variables | Degrees of freedom | Al F (P) | Ca F (P) | Cu F (P) | Fe F (P) | Mg F (P) | Mn F (P) | P F (P) | K F (P) | Na F (P) | Zn F (P) |
|---|---|---|---|---|---|---|---|---|---|---|---|---|
| **Hyphal growth** | Protein | 1 | 146.3 (<2.2e-16) *** | 204.51 (<2.2e-16) *** | 215.05 (<2.2e-16) *** | 184.31 (<2.2e-16) *** | 309.44 (<2.2e-16) *** | 171.16 (<2.2e-16) *** | 91.88 (<2.2e-16) *** | 292.39 (<2.2e-16) *** | 299.05 (<2.2e-16) *** | 220.07 (<2.2e-16) *** |
| | Carbohydrates | 1 | 260.5 (<2.2e-16) *** | 251.07 (<2.2e-16) *** | 251.47 (<2.2e-16) *** | 193.22 (<2.2e-16) *** | 301.47 (<2.2e-16) *** | 333.98 (<2.2e-16) *** | 157.23 (<2.2e-16) *** | 287.58 (<2.2e-16) *** | 276.4 (<2.2e-16) *** | 220.37 (<2.2e-16) *** |
| | Mineral | 1 | 183.6 (<2.2e-16) *** | 756.24 (<2.2e-16) *** | 206.29 (<2.2e-16) *** | 123.85 (<2.2e-16) *** | 586.01 (<2.2e-16) *** | 231.5 (<2.2e-16) *** | 149.31 (<2.2e-16) *** | 668.34 (<2.2e-16) *** | 786.29 (<2.2e-16) *** | 1043.3 (<2.2e-16) *** |
| | Protein$^2$ | 1 | 31.8 (2.4e-08) *** | 12.23 (5e-04) *** | 42.87 (1.01e-10) *** | 20.37 (7.4e-06) *** | 9.29 (2.4e-03) ** | 20.83 (5.86e-06) *** | 50.63 (2.63e-12) *** | 17.15 (3.9e-05) *** | 10.03 (1.6e-03) ** | 29.18 (8.9e-08) *** |
| | Carbohydrates$^2$ | 1 | 114.4 (<2.2e-16) *** | 111.77 (<2.2e-16) *** | 57.42 (1.05e-13) *** | 112.62 (<2.2e-16) *** | 157.29 (<2.2e-16) *** | 130.63 (<2.2e-16) *** | 58.72 (5.68e-14) *** | 179.33 (<2.2e-16) *** | 164.62 (<2.2e-16) *** | 118.24 (<2.2e-16) *** |
| | Mineral$^2$ | 1 | 4.5 (0.033) * | 127.99 (<2.2e-16) *** | 15.04 (1.15e-4) *** | 44.9 (4.1e-11) *** | 131.49 (<2.2e-16) *** | 0.96 (0.33) | 1.04 (0.31) | 1.83 (0.18) | 3.59 (0.06) | 75.1 (<2.2e-16) *** |
| | Protein x Carbohydrates | 1 | 59.01 (4.9e-14) *** | 81.51 (<2.2e-16) *** | 75.7 (<2.2e-16) *** | 70.18 (2.68e-16) *** | 111.19 (<2.2e-16) *** | 49.4 (4.73e-12) *** | 68.41 (6.1e-16) *** | 115.08 (<2.2e-16) *** | 103.09 (<2.2e-16) *** | 108.41 (<2.2e-16) *** |
| | Protein x Mineral | 1 | 248.8 (<2.2e-16) *** | 47.49 (1.18e-11) *** | 19.22 (1.33e-05) *** | 159.25 (<2.2e-16) *** | 9.98 (1.7e-03) ** | 59.75 (3.49e-14) *** | 166.5 (<2.2e-16) *** | 14.37 (1.6e-04) *** | 22 (3.24e-06) *** | 50.38 (3e-12) *** |
| | Carbohydrates x Mineral | 1 | 18.9 (1.6e-05) *** | 30.91 (3.77e-08) *** | 1.89 (0.17) | 43.68 (7.35e-11) *** | 6.77 (9.4e-03) ** | 2.9 (0.09) | 47.01 (1.49e-11) *** | 35.95 (3.2e-09) *** | 27.04 (2.58e-07) *** | 60.2 (2.8e-14) *** |
| | Protein x Carbohydrtaes. x Mineral | 1 | 13.7 (2.3e-04) *** | 1.29 (0.26) | 0.46 (0.50) | 6.88 (8.9e-03) ** | 2.39 (0.12) | 26.66 (3.11e-07) *** | 1.03 (0.31) | 2.18 (0.14) | 1.57 (0.21) | 1.34 (0.25) |
| | Residuals | 745 | | | | | | | | | | |

Mapping hyphal growth across nutritional landscape varying across 7 Pr:C ratios (9:1, 6:1, 3:1, 1:1, 1:3, 1:6, 1:9 Pr:C), 3 Pr+C dilutions (4, 8, and 25 g/L) and 3 mineral concentrations for each mineral (baseline without mineral + 2 mineral concentrations (see methods)). For each mineral we prepared 42 diets (7 Pr:C * 3 Pr+C * 2 minerals concentrations = 42 diets) for a total of 441 diets ((42 * 10 minerals) + 21 baseline diets). Shown are the results of least-square regressions estimating parametric nonlinear response surfaces between control and each mineral element individually using linear and quadratic components of protein, carbohydrate, and mineral concentrations (Mineral) in diets and their interactions. F indicates F-value and P the probability of F. Significance of estimated coefficients: *P < 0.05, **P < 0.01, ***P < 0.001.

**Table S4: Significance tests for univariate models for hyphal growth in presence of mineral elements.**

| Model – Hyphal growth | Al | Ca | Cu | Fe | Mg | Mn | P | K | Na | Zn |
|---|---|---|---|---|---|---|---|---|---|---|
| **Degrees of freedom model** | 10 | 10 | 10 | 10 | 10 | 10 | 10 | 10 | 10 | 10 |
| **Degrees of freedom residuals** | 745 | 745 | 745 | 745 | 745 | 745 | 745 | 745 | 745 | 745 |
| **Adjusted $R^2$** | 0.5867 | 0.6814 | 0.5369 | 0.557 | 0.6815 | 0.5741 | 0.5087 | 0.68 | 0.6904 | 0.7174 |
| **Mean Square Model Residuals** | 22640 | 21832 | 26532 | 26573 | 20986 | 25767 | 30531 | 20861 | 20217 | 21367 |
| **F-statistic** | 108.2 | 162.5 | 88.54 | 95.93 | 162.5 | 102.8 | 79.18 | 161.4 | 169.4 | 192.7 |
| **P-value** | <2.2e-16*** | <2.2e-16*** | <2.2e-16*** | <2.2e-16*** | <2.2e-16*** | <2.2e-16*** | <2.2e-16*** | <2.2e-16*** | <2.2e-16*** | <2.2e-16*** |

Mapping hyphal growth across nutritional landscape varying across 7 Pr:C ratios (9:1, 6:1, 3:1, 1:1, 1:3, 1:6, 1:9 Pr:C), 3 Pr+C dilutions (4, 8, and 25 g/L) and 3 mineral concentrations for each mineral (baseline without mineral + 2 mineral concentrations (see methods)). For each mineral we prepared 42 diets (7 Pr:C * 3 Pr+C * 2 minerals concentrations = 42 diets) for a total of 441 diets ((42 * 10 minerals) + 21 baseline diets). Shown are the characteristics and significance of models for each mineral element individually from Table S12. Significance of estimated coefficients: *P < 0.05, **P < 0.01, ***P < 0.001.

**Table S5: Estimated coefficients for hyphal growth in presence of mineral elements.**

| Response variable | Explanatory variables | Al β (SE) | Ca β (SE) | Cu β (SE) | Fe β (SE) | Mg β (SE) | Mn β (SE) | P β (SE) | K β (SE) | Na β (SE) | Zn β (SE) |
|---|---|---|---|---|---|---|---|---|---|---|---|
| **Hyphal growth** | Protein | 49.45 (6.01)*** | 46.56 (5.9)*** | 59.58 (6.52)*** | 52.42 (6.51)*** | 50.72 (5.78)*** | 48.34 (6.42) *** | 66.14 (6.99)*** | 55.78 (5.76)*** | 48.52 (5.68)*** | 58.16 (5.85)*** |
| | Carbohydrates | 89.26 (6.01)*** | 91.93 (5.9)*** | 86.95 (6.51)*** | 98.13 (6.51)*** | 103.89 (5.78)*** | 95.5 (6.42)*** | 91.11 (6.99)*** | 107.58 (5.76)*** | 101.27 (5.67)*** | 98.28 (5.85)*** |
| | Mineral | -1464 (922.4) | -2870.9 (181.15)*** | -22700 (3102)*** | 4230 (833.3)*** | -243.89 (17.91)*** | -15090 (3057)*** | -858.98 (332.76) * | -213.66 (128.87) | -475.56 (134.2)*** | -41190 (2784)*** |
| | $Protein^2$ | -2.27 (0.21)*** | -2.09 (0.21)*** | -2.66 (0.23)*** | -2.37 (0.23)*** | -2.24 (0.2)*** | -2.17 (0.23)*** | -2.89 (0.24)*** | -2.44 (0.2)*** | -2.17 (0.2)*** | -2.55 (0.2)*** |
| | $Carbohydrates^2$ | -2.76 (0.21)*** | -2.88 (0.21)*** | -2.62 (0.23)*** | -3.08 (0.23)*** | -3.32 (0.2)*** | -2.97 (0.23)*** | -2.75 (0.24)*** | -3.46 (0.2)*** | -3.25 (0.2)*** | -3.08 (0.2)*** |
| | $Mineral^2$ | -1580 (7426)* | 3300.02 (291.7)*** | 399400 (103000)*** | -22830 (3407)*** | 16.87 (1.47)*** | 99340 (101500) | -1128.51 (1104.78) | 14.5 (10.73) | 105.02 (55.46) | 800900 (92420)*** |
| | Protein x Carbohydrates | -3.51 (0.42)*** | -3.52 (0.41)*** | -3.77 (0.46)*** | -3.97 (0.45)*** | -4.02 (0.4)*** | -3.68 (0.45)*** | -4.06 (0.49)*** | -4.1 (0.4)*** | -3.82 (0.39)*** | -4.01 (0.41)*** |
| | Protein x Mineral | 248.5 (43.53)*** | 39.61 (8.55)*** | 527.1 (176.6)** | 113 (23.87)*** | 1.26 (0.42)** | 30.02 (174) | 117.65 (18.94)*** | 1.33 (0.42)** | 7.29 (2.08)*** | 743.1 (158.5)*** |
| | Carbohydrates x Mineral | -239.7 (43.52)*** | -18.91 (8.55)* | -39.2 (176.5) | -140.9 (23.86)*** | -0.09 (0.42) | -572.7 (174)** | -89.37 (18.94)*** | -0.92 (0.42)* | -3.98 (2.08) | -544.8 (158.4)*** |
| | Protein x Carbohydrates x Mineral | 29.3 (7.92)*** | -1.77 (1.56) | -21.7 (32.13) | 11.39 (4.34)** | -0.12 (0.08) | 163.5 (31.67)*** | 3.5 (3.45) | -0.11 (0.08) | -0.48 (0.38) | -33.4 (28.84) |

Mapping hyphal growth across nutritional landscape varying across 7 Pr:C ratios (9:1, 6:1, 3:1, 1:1, 1:3, 1:6, 1:9 Pr:C), 3 Pr+C dilutions (4, 8, and 25 g/L) and 3 mineral concentrations for each mineral (baseline without mineral + 2 mineral concentrations (see methods)). For each mineral we prepared 42 diets (7 Pr:C * 3 Pr+C * 2 minerals concentrations = 42 diets) for a total of 441 diets ((42 * 10 minerals) + 21 baseline diets). Shown are the results of least-square regressions estimating parametric nonlinear response surfaces between control and each mineral element individually using linear and quadratic components of protein, carbohydrate, and mineral concentrations (Mineral) composition in diets and their interactions. β indicates coefficient of the model for each parameter with SE that indicates standard error. Significance of estimated coefficients: *$P < 0.05$, **$P < 0.01$, ***$P < 0.001$.

**Table S6: Location of the *A. colombica* colonies used in this study conducted in Sobéranía Park (Panama).**

| ***Species*** | **Colony ID** | **GPS coordinates** | **Location** |
|---|---|---|---|
| *Atta colombica* | Colony 1 | 9.15089, -79.73305 | Pipeline road |
| | Colony 2 | 9.15471, -79.73512 | Pipeline road |
| | Colony 3 | 9.11929, -79.69494 | Laguna forest |
| | Colony 4 | 9.11672, -79.69514 | Laguna forest |
| | Colony 5 | 9.10544, -79.61901 | Camino de cruces |
| | Colony 6 | 9.10307, -79.61641 | Camino de cruces |

**Table S7: Plant species identification from NCBI Blast n results with E-value, percent identity and NCBI accession numbers for each sample.**

| Field sample ID | Identification | E-value | Per. Ident. | Accession number | Field sample ID | Identification | E-value | Per. Ident. | Accession number | Field sample ID | Identification | E-value | Per. Ident. | Accession number |
|---|---|---|---|---|---|---|---|---|---|---|---|---|---|---|
| **sp65** | *Aegiphila monstrosa* | 9.00E-112 | 95.42% | MT509077 | **sp7** | *Erythrina fusca* | 5.00E-109 | 92.81% | MT509032 | **sp49** | *Mangifera indica* | 1.00E-176 | 100% | MT509061 |
| **sp71** | *Adelia triloba* | 1.00E-79 | 98.84% | MT509083 | **bcc8** | *Ficus adhatodifolia* | 2.00E-112 | 100% | MT509022 | **sp29** | *Mangifera indica* | 1.00E-74 | 100% | MT509045 |
| **sp73** | *Alchornea davidii* | 2.00E-117 | 95.88% | MT509085 | **sp56** | *Ficus adhatodifolia* | 2.00E-132 | 98.21% | MT509068 | **sc18** | *Mangifera indica* | 2.00E-173 | 99.42% | MT509026 |
| **sp26** | *Amorimia camporum* | 3.00E-86 | 89.71% | MT509042 | **sp55** | *Ficus adhatodifolia* | 1.00E-125 | 100% | MT509067 | **sp47** | *Mangifera indica* | 1.00E-171 | 100% | MT509060 |
| **sp45** | *Amorimia camporum* | 4.00E-95 | 93.15% | MT509058 | **sp64** | *Ficus adhatodifolia* | 1.00E-130 | 99.62% | MT509076 | **sp33** | *Mangifera sp.* | 5.00E-98 | 99.02% | MT509048 |
| **sc8c12** | *Astrocaryum standleyanum* | 9.00E-168 | 97.71% | MT509024 | **sp5** | *Ficus calimana* | 2.00E-99 | 93.65% | MT509031 | **sp88** | *Miconia argentea* | 2.00E-91 | 99.48% | MT509099 |
| **sp54** | *Astrocaryum standleyanum* | 8.00E-102 | 96.93% | MT509066 | **sp46** | *Ficus sp.* | 3.00E-105 | 98.64% | MT509059 | **sp20** | *Miconia argentea* | 2.00E-168 | 99.70% | MT509039 |
| **sp77** | *Bignonia capreolata* | 4.00E-110 | 92.07% | MT509089 | **sp58** | *Ficus tonduzii* | 1.00E-156 | 98.74% | MT509070 | **sp72** | *Microcos paniculata* | 9.00E-57 | 82.67% | MT509084 |
| **sp51** | *Bignonia capreolata* | 2.00E-103 | 91.76% | MT509063 | **sp53** | *Ficus tonduzii* | 5.00E-155 | 97.84% | MT509065 | **sp81** | *Odontadenia glauca* | 2.00E-29 | 77.03% | MT509093 |
| **sp32** | *Bougainvillea spectabilis* | 4.00E-135 | 98.92% | MT509047 | **sp97** | *Ficus tonduzii* | 2.00E-163 | 99.69% | MT509108 | **sp95** | *Odontadenia glauca* | 5.00E-48 | 87.29% | MT509106 |
| **sp14** | *Brownea sp.* | 3.00E-151 | 97.78% | MT509035 | **sc2** | *Ficus tonduzii* | 2.00E-164 | 99.69% | MT509023 | **sp79** | *Pinzona coriacea* | 2.00E-83 | 90.87% | MT509091 |
| **sp63** | *Bucida buceras* | 1.00E-105 | 91.90% | MT509075 | **sc12** | *Ficus tonduzii* | 2.00E-164 | 99.69% | MT509025 | **sp75** | *Pithecoctenium sp.* | 5.00E-105 | 89.71% | MT509087 |
| **sp66** | *Cissus quadrangularis* | 1.00E-32 | 96.74% | MT509078 | **sp27** | *Hieronyma alchorneoides* | 3.00E-146 | 99.65% | MT509043 | **sp60** | *Pithecoctenium sp.* | 2.00E-83 | 88.15% | MT509072 |
| **sp21** | *Combretum coccineum* | 6.00E-99 | 90.10% | MT509040 | **sp57** | *Hieronyma alchorneoides* | 4.00E-135 | 100% | MT509069 | **sp2** | *Poulsenia armata* | 4.00E-88 | 100% | MT509028 |
| **sp37** | *Combretum coccineum* | 2.00E-98 | 90.78% | MT509052 | **sp28** | *Hieronyma alchorneoides* | 7.00E-143 | 98.97% | MT509044 | **sp44** | *Rinorea hummelii* | 7.00E-93 | 90.60% | MT509057 |
| **sp41** | *Combretum coccineum* | 2.00E-93 | 89.96% | MT509055 | **sp96** | *Hieronyma alchorneoides* | 1.00E-130 | 99.62% | MT509107 | **sp86** | *Rinorea hummelii* | 6.00E-109 | 93.43% | MT509098 |
| **sp38** | *Combretum coccineum* | 5.00E-115 | 91.43% | MT509053 | **sp42** | *Hylenaea praecelsa* | 1.00E-104 | 100% | MT509056 | **sp18** | *Serjania mexicana* | 4.00E-155 | 99.04% | MT509037 |
| **sp61** | *Combretum coccineum* | 2.00E-103 | 90.82% | MT509073 | **sp78** | *Hylenaea praecelsa* | 1.00E-151 | 100% | MT509090 | **sp82** | *Swartzia guatemalensis* | 2.00E-134 | 98.91% | MT509094 |
| **sp62** | *Combretum coccineum* | 9.00E-102 | 90.48% | MT509074 | **sp74** | *Hylenaea praecelsa* | 3.00E-100 | 99.51% | MT509086 | **sp76** | *Swartzia guatemalensis* | 1.00E-136 | 98.92% | MT509088 |
| **sp25** | *Combretum coccineum* | 2.00E-68 | 89.19% | MT509041 | **sp85** | *Hylenaea praecelsa* | 1.00E-151 | 100% | MT509097 | **sp1** | *Swietenia macrophylla* | 1.00E-171 | 99.70% | MT509027 |
| **sp12** | *Croton spruceanus* | 2.00E-153 | 98.41% | MT509033 | **sp84** | *Inga laurina* | 2.00E-128 | 99.61% | MT509096 | **sp17** | *Swietenia macrophylla* | 2.00E-163 | 98.21% | MT509036 |
| **sp94** | *Davilla nitida* | 4.00E-130 | 99.62% | MT509105 | **sp4** | *Iseia luxurians* | 5.00E-79 | 99.41% | MT509030 | **sp13** | *Terminalia hainanensis* | 5.00E-94 | 92.13% | MT509034 |
| **sp19** | *Davilla nitida* | 4.00E-130 | 98.16% | MT509038 | **sp69** | *Luehea divaricata* | 1.00E-43 | 94.26% | MT509081 | **sp36** | *Terminalia hainanensis* | 1.00E-95 | 91.60% | MT509051 |
| **sp70** | *Davilla nitida* | 4.00E-78 | 99.40% | MT509082 | **sp80** | *Luehea divaricata* | 4.00E-85 | 91.25% | MT509092 | **sp68** | *Terminalia hainanensis* | 5.00E-94 | 92.13% | MT509080 |
| **sp89** | *Davilla nitida* | 1.00E-130 | 99.62% | MT509100 | **sp90** | *Luehea divaricata* | 3.00E-92 | 90.91% | MT509101 | **sp40** | *Terminalia pallida* | 4.00E-105 | 92.20% | MT509054 |
| **sp92** | *Davilla nitida* | 5.00E-139 | 100% | MT509103 | **sp3** | *Ludwigia abyssinica* | 6.00E-128 | 98.86% | MT509029 | **sp34** | *Trophis racemosa* | 3.00E-75 | 98.78% | MT509049 |
| **sp35** | *Doliocarpus dentatus* | 1.00E-77 | 97.71% | MT509050 | **sp50** | *Lupinus angustifolius* | 7.00E-20 | 98.39% | MT509062 | **sp52** | *Trophis racemosa* | 1.00E-121 | 98.43% | MT509064 |
| **sp91** | *Dolichandra unguis-cati* | 6.00E-99 | 89.80% | MT509102 | **sp83** | *Mabea sp.* | 1.00E-131 | 97.50% | MT509095 | **sp59** | *Vitex megapotamica* | 3.00E-80 | 94.12% | MT509071 |
| **sp93** | *Dolichandra unguis-cati* | 2.00E-99 | 90.03% | MT509104 | **sp31** | *Mangifera indica* | 6.00E-123 | 100% | MT509046 | **sp67** | *Vitex megapotamica* | 1.00E-73 | 96.05% | MT509079 |

**Table S8: Nutrient and mineral composition of plant species per substrate type harvested by *A. colombica*.**

| Plant species (*: leaves/**:flowers/***:fruits) | TNC % | Protein % | Al (ug/g) | Cu (ug/g) | Fe (ug/g) | Mn (ug/g) | Na (ug/g) | Zn (ug/g) | Ca (mg/g) | K (mg/g) | Mg (mg/g) | P (mg/g) | Colony ID | Dry mass (g) |
|---|---|---|---|---|---|---|---|---|---|---|---|---|---|---|
| ***Aegiphila monstrosa* (*)** | 17.6 | 14.4 | 17.1 | 24.5 | 93.5 | 37.4 | 63.7 | 89.6 | 5.7 | 14.2 | 3.1 | 1.8 | colony 6 | 0.04 |
| ***Adelia triloba* (*)** | 15.2 | 17.6 | 21.5 | 10.2 | 54.3 | 72.3 | 88.6 | 158.1 | 15.2 | 13.8 | 4.1 | 1.9 | colony 1 | 0.10 |
| ***Alchornea davidii* (*)** | 12.4 | 14.4 | 36.4 | 14.2 | 78.8 | 40.7 | 149.1 | 23.1 | 4.5 | 8.0 | 2.7 | 2.3 | colony 6 | 2.25 |
| ***Amorimia camporum* (*)** | 3.0 | 16.1 | 56.8 | 15.7 | 70.3 | 20.0 | 81.4 | 60.1 | 5.1 | 16.6 | 2.3 | 2.7 | colony 6 | 5.61 |
| ***Amorimia camporum* (*)** | 15.5 | 11.3 | 1031.1 | 16.5 | 51.3 | 22.6 | 95.5 | 33.6 | 6.2 | 16.6 | 2.6 | 1.6 | colony 2 | 0.04 |
| ***Astrocaryum standleyanum* (***)** | 33.1 | 5.3 | 13.5 | 15.4 | 29.2 | 8.9 | 51.6 | 29.0 | 0.8 | 18.2 | 1.0 | 1.0 | colony 4 | 0.60 |
| ***Astrocaryum standleyanum* (***)** | 19.3 | 6.1 | 132.9 | 8.6 | 142.5 | 8.8 | 99.8 | 22.0 | 0.7 | 21.8 | 1.2 | 0.9 | colony 3 | 0.11 |
| ***Bignonia capreolata* (*)** | 17.3 | 20.7 | 114.1 | 13.8 | 136.7 | 16.5 | 24.8 | 20.8 | 2.7 | 9.6 | 1.6 | 2.1 | colony 2 | 0.23 |
| ***Bignonia capreolata* (*)** | 8.4 | 15.0 | 97.4 | 27.2 | 130.7 | 16.6 | 64.7 | 26.5 | 2.9 | 16.3 | 1.2 | 2.0 | colony 4 | 1.02 |
| ***Bougainvillea spectabilis* (*)** | 15.6 | 34.8 | 110.8 | 7.5 | 183.5 | 56.4 | 624.0 | 13.8 | 33.0 | 36.2 | 6.7 | 2.1 | colony 4 | 0.04 |
| ***Brownea sp.* (**)** | 25.8 | 3.2 | 230.3 | 27.7 | 309.5 | 18.4 | 2488.1 | 118.9 | 3.5 | 80.9 | 14.9 | 12.0 | colony 3 | 0.03 |
| ***Bucida buceras* (**)** | 10.5 | 11.9 | 52.0 | 17.4 | 59.6 | 56.4 | 439.7 | 26.6 | 5.0 | 23.5 | 2.4 | 1.8 | colony 6 | 0.84 |
| ***Cissus quadrangularis* (*)** | 23.0 | 21.1 | 46.8 | 13.1 | 82.6 | 25.1 | 34.2 | 32.8 | 1.5 | 18.2 | 1.9 | 2.9 | colony 6 | 0.04 |
| ***Combretum coccineum* (*)** | 23.0 | 21.8 | 126.9 | 23.7 | 152.0 | 24.7 | 77.3 | 46.3 | 3.2 | 20.7 | 2.0 | 4.1 | colony 5 | 5.98 |
| ***Combretum coccineum* (*)** | 6.8 | 22.3 | 51.2 | 25.2 | 96.0 | 34.9 | 73.0 | 53.7 | 6.7 | 23.8 | 2.5 | 4.7 | colony 5 | 10.09 |
| ***Combretum coccineum* (*)** | 13.7 | 15.0 | 39.1 | 17.4 | 59.6 | 83.6 | 60.7 | 54.5 | 9.0 | 14.4 | 2.2 | 2.3 | colony 2 | 1.95 |
| ***Combretum coccineum* (*)** | 11.1 | 28.7 | 46.3 | 34.9 | 108.9 | 23.5 | 76.8 | 68.2 | 1.7 | 25.0 | 2.4 | 6.4 | colony 5 | 0.06 |
| ***Combretum coccineum* (*)** | 10.5 | 18.3 | 39.6 | 14.5 | 236.2 | 160.1 | 140.7 | 110.7 | 18.8 | 12.6 | 3.8 | 1.5 | colony 6 | 0.27 |
| ***Combretum coccineum* (*)** | 12.0 | 17.4 | 43.7 | 20.9 | 64.6 | 62.9 | 129.3 | 54.3 | 6.5 | 19.2 | 2.3 | 2.4 | colony 6 | 0.08 |
| ***Combretum coccineum* (*)** | 8.0 | 17.3 | 49.0 | 21.3 | 71.6 | 35.0 | 67.9 | 42.3 | 2.9 | 18.9 | 2.0 | 3.0 | colony 6 | 5.34 |
| ***Croton spruceanus* (*)** | 16.8 | 29.7 | 46.5 | 11.7 | 119.5 | 224.9 | 1246.4 | 40.0 | 15.9 | 13.4 | 7.4 | 1.7 | colony 3 | 5.10 |
| ***Davilla nitida* (*)** | 15.5 | 8.0 | 85.7 | 8.8 | 86.8 | 128.4 | 187.9 | 10.7 | 3.8 | 10.0 | 3.3 | 0.8 | colony 1 | 0.08 |
| ***Davilla nitida* (*)** | 13.0 | 14.0 | 31.9 | 9.0 | 50.0 | 235.3 | 67.5 | 14.0 | 9.5 | 13.4 | 3.0 | 1.3 | colony 3 | 0.01 |
| ***Davilla nitida* (*)** | 17.4 | 7.3 | 23.4 | 7.6 | 33.8 | 137.4 | 211.1 | 8.3 | 4.3 | 7.7 | 3.3 | 0.9 | colony 1 | 0.62 |
| ***Davilla nitida* (*)** | 12.0 | 8.3 | 148.8 | 6.6 | 161.1 | 38.3 | 90.3 | 19.2 | 4.0 | 12.0 | 2.5 | 1.3 | colony 1 | 0.99 |
| ***Davilla nitida* (*)** | 17.5 | 5.1 | 102.8 | 9.8 | 115.4 | 105.3 | 298.4 | 11.7 | 3.2 | 10.9 | 3.3 | 1.1 | colony 1 | 0.05 |

| | | | | | | | | | | | | | |
|---|---|---|---|---|---|---|---|---|---|---|---|---|---|
| ***Doliocarpus dentatus* (*)** | 11.8 | 13.0 | 99.9 | 12.3 | 114.8 | 77.7 | 164.6 | 18.4 | 7.2 | 10.5 | 3.6 | 1.4 | colony 3 | 1.13 |
| ***Dolichandra unguis-cati* (*)** | 17.0 | 21.9 | 232.4 | 15.4 | 259.2 | 149.3 | 87.4 | 22.5 | 12.7 | 11.1 | 4.0 | 1.9 | colony 1 | 0.22 |
| ***Dolichandra unguis-cati* (*)** | 20.6 | 16.4 | 134.8 | 11.8 | 99.7 | 34.4 | 119.7 | 17.2 | 2.4 | 13.1 | 2.7 | 1.5 | colony 1 | 0.10 |
| ***Erythrina fusca* (*)** | 13.6 | 20.8 | 138.0 | 13.5 | 200.5 | 580.2 | 1070.4 | 30.3 | 10.8 | 28.6 | 5.8 | 2.3 | colony 4 | 0.02 |
| ***Ficus adhatodifolia* (**)** | 12.4 | 8.8 | 44.5 | 11.7 | 52.0 | 74.7 | 79.4 | 11.1 | 4.4 | 20.7 | 1.7 | 1.0 | colony 3 | 0.41 |
| ***Ficus adhatodifolia* (*)** | 5.3 | 4.2 | 88.4 | 7.3 | 36.5 | 55.1 | 102.7 | 11.1 | 20.5 | 15.6 | 4.0 | 0.7 | colony 5 | 3.44 |
| ***Ficus adhatodifolia* (***)** | 32.1 | -1.1(ǂ) | 644.1 | 7.4 | 610.7 | 32.5 | 210.9 | 15.1 | 9.2 | 16.5 | 2.4 | 0.8 | colony 5 | 0.80 |
| ***Ficus adhatodifolia* (***)** | 17.5 | 4.4 | 91.5 | 3.8 | 55.3 | 22.9 | 246.0 | 9.2 | 26.2 | 13.6 | 2.7 | 0.7 | colony 6 | 0.18 |
| ***Ficus calimana* (*)** | 18.8 | 6.9 | 90.0 | 7.2 | 137.8 | 29.3 | 257.3 | 19.1 | 14.8 | 17.3 | 2.8 | 1.3 | colony 4 | 0.41 |
| ***Ficus sp.* (*)** | 30.0 | 16.4 | 62.1 | 10.1 | 105.9 | 23.6 | 49.2 | 36.7 | 10.7 | 19.5 | 3.7 | 2.8 | colony 4 | 14.06 |
| ***Ficus tonduzii* (*)** | 18.5 | 23.9 | 276.6 | 10.7 | 54.8 | 45.6 | 80.4 | 19.0 | 9.9 | 17.9 | 3.3 | 2.5 | colony 5 | 0.17 |
| ***Ficus tonduzii* (*)** | 4.7 | 6.3 | 161.3 | 6.9 | 112.0 | 70.5 | 39.0 | 15.6 | 12.2 | 17.6 | 5.4 | 1.1 | colony 3 | 4.02 |
| ***Ficus tonduzii* (***)** | 19.7 | 17.1 | 107.5 | 8.1 | 63.2 | 187.5 | 41.9 | 31.0 | 30.5 | 19.1 | 8.1 | 3.0 | colony 1 | 0.80 |
| ***Ficus tonduzii* (***)** | 26.3 | 0.5 | 164.5 | 8.5 | 66.9 | 37.4 | 44.8 | 9.5 | 5.6 | 12.9 | 2.0 | 0.9 | colony 1 | 0.54 |
| ***Ficus tonduzii* (***)** | 31.5 | 4.6 | 87.5 | 6.6 | 24.9 | 16.5 | 56.2 | 9.2 | 7.0 | 23.6 | 2.4 | 1.2 | colony 5 | 0.13 |
| ***Hieronyma alchorneoides* (*)** | 13.2 | 12.8 | 59.2 | 8.1 | 64.6 | 18.0 | 46.4 | 14.7 | 3.7 | 9.5 | 2.5 | 2.0 | colony 6 | 6.23 |
| ***Hieronyma alchorneoides* (*)** | 13.1 | 12.6 | 18.9 | 9.3 | 45.3 | 16.4 | 18.3 | 13.9 | 4.1 | 10.3 | 2.2 | 2.3 | colony 5 | 9.09 |
| ***Hieronyma alchorneoides* (***)** | 45.2 | 3.5 | 768.4 | 5.7 | 827.0 | 21.7 | 200.0 | 8.7 | 1.6 | 4.8 | 1.0 | 0.9 | colony 6 | 1.34 |
| ***Hieronyma alchorneoides* (***)** | 38.3 | 10.4 | 107.3 | 7.0 | 102.7 | 9.5 | 117.8 | 10.1 | 2.5 | 9.6 | 1.2 | 1.2 | colony 6 | 0.44 |
| ***Hylenaea praecelsa* (*)** | 7.0 | 16.8 | 119.6 | 11.3 | 98.5 | 26.7 | 27.0 | 26.6 | 6.3 | 15.0 | 1.8 | 2.4 | colony 2 | 5.42 |
| ***Hylenaea praecelsa* (*)** | 15.2 | 22.4 | 253.8 | 10.2 | 266.9 | 36.5 | 57.9 | 65.1 | 9.9 | 19.2 | 4.6 | 1.9 | colony 2 | 0.08 |
| ***Hylenaea praecelsa* (*)** | 5.3 | 11.7 | 45.9 | 8.8 | 62.0 | 30.0 | 33.8 | 24.0 | 14.7 | 9.4 | 2.2 | 1.5 | colony 2 | 5.84 |
| ***Hylenaea praecelsa* (*)** | 15.0 | 15.8 | 23.4 | 10.7 | 45.8 | 38.0 | 195.0 | 24.4 | 17.0 | 10.2 | 1.8 | 1.7 | colony 2 | 0.28 |
| ***Inga laurina* (**)** | 19.2 | 19.2 | 89.4 | 11.7 | 125.5 | 28.4 | 57.2 | 34.2 | 1.6 | 12.2 | 1.1 | 2.4 | colony 2 | 0.31 |
| ***Iseia luxurians* (*)** | 12.4 | 15.8 | 27.6 | 18.7 | 51.6 | 17.0 | 97.7 | 18.4 | 3.5 | 13.8 | 1.1 | 1.6 | colony 4 | 5.13 |
| ***Luehea divaricata* (*)** | 4.8 | 14.4 | 22.1 | 21.6 | 44.4 | 202.4 | 14.0 | 21.8 | 4.2 | 15.8 | 3.1 | 3.6 | colony 1 | 16.66 |
| ***Luehea divaricata* (*)** | 15.8 | 14.1 | 24.5 | 21.8 | 49.8 | 181.8 | 37.8 | 20.8 | 3.3 | 13.3 | 2.8 | 3.0 | colony 1 | 9.89 |
| ***Luehea divaricata* (*)** | 12.7 | 11.5 | 83.7 | 21.4 | 110.6 | 167.6 | 51.6 | 21.3 | 3.3 | 14.6 | 2.9 | 3.4 | colony 1 | 1.07 |
| ***Ludwigia abyssinica* (*)** | 11.9 | 23.0 | 123.2 | 16.2 | 269.7 | 316.5 | 179.7 | 54.4 | 13.8 | 13.9 | 4.1 | 3.8 | colony 4 | 0.76 |

| | | | | | | | | | | | | | |
|---|---|---|---|---|---|---|---|---|---|---|---|---|---|
| ***Lupinus angustifolius* (*)** | 17.4 | 23.0 | 74.5 | 5.6 | 103.6 | NA | 143.5 | NA | 23.0 | 5.8 | 3.6 | 0.9 | colony 4 | 1.52 |
| ***Mabea sp.* (*)** | 11.8 | 15.9 | 44.7 | 10.8 | 87.1 | 114.9 | 241.6 | 27.0 | 8.6 | 4.0 | 2.4 | 1.0 | colony 2 | 0.31 |
| ***Mangifera indica* (*)** | 15.2 | 10.3 | 84.2 | 8.2 | 111.1 | 46.6 | 49.0 | 15.3 | 4.1 | 11.3 | 2.1 | 2.1 | colony 4 | 2.15 |
| ***Mangifera indica* (*)** | 17.1 | 12.9 | 28.1 | 8.6 | 42.6 | 89.4 | 148.9 | 16.7 | 16.0 | 5.3 | 3.7 | 1.1 | colony 4 | 1.66 |
| ***Mangifera indica* (*)** | 8.3 | 28.5 | 639.1 | 12.7 | 783.8 | 43.3 | 164.6 | 51.3 | 3.1 | 23.4 | 2.7 | 4.7 | colony 4 | 7.85 |
| ***Mangifera indica* (***)** | 44.3 | 2.4 | 202.1 | 4.2 | 177.6 | 26.4 | 33.2 | 5.2 | 1.3 | 8.8 | 1.2 | 0.6 | colony 4 | 0.13 |
| ***Mangifera indica* (***)** | 23.5 | 4.6 | 4175.0 | 15.2 | NA | 196.6 | 152.9 | 24.3 | 3.5 | 8.6 | 3.1 | 0.8 | colony 4 | 0.19 |
| ***Mangifera sp.* (*)** | 18.0 | 12.1 | 66.2 | 10.8 | 75.7 | 52.1 | 42.6 | 22.2 | 2.6 | 12.5 | 1.8 | 3.0 | colony 4 | 0.06 |
| ***Miconia argentea* (*)** | 10.7 | 15.6 | NA | 8.2 | 83.8 | 76.0 | 469.0 | 37.8 | 9.3 | 4.3 | 3.4 | 1.4 | colony 1 | 2.49 |
| ***Miconia argentea* (***)** | 24.5 | 17.5 | 9468.7 | 7.2 | 57.5 | 52.9 | 171.5 | 42.3 | 9.9 | 8.1 | 2.5 | 1.5 | colony 3 | 0.07 |
| ***Microcos paniculata* (*)** | 13.6 | 18.2 | 32.9 | 12.1 | 74.0 | 32.5 | 22.4 | 26.4 | 4.4 | 16.9 | 2.5 | 1.9 | colony 6 | 12.99 |
| ***Odontadenia glauca* (*)** | 19.4 | 20.9 | 35.9 | 11.8 | 69.9 | 730.4 | 58.5 | NA | 8.9 | 13.6 | 2.8 | 2.0 | colony 1 | 0.42 |
| ***Odontadenia glauca* (**)** | 21.1 | 9.7 | 58.3 | 11.0 | 64.4 | 113.2 | 30.1 | 12.4 | 2.1 | 9.9 | 1.3 | 1.2 | colony 6 | 0.24 |
| ***Pinzona coriacea* (*)** | 4.5 | 10.8 | 30.8 | 7.9 | 56.7 | 36.5 | 61.2 | 18.2 | 4.5 | 11.6 | 2.6 | 1.4 | colony 1 | 5.39 |
| ***Pithecoctenium sp.* (*)** | 14.8 | 23.6 | 262.2 | 29.7 | 358.4 | 35.0 | 64.4 | 67.7 | 4.2 | 21.3 | 2.4 | 3.8 | colony 2 | 1.32 |
| ***Pithecoctenium sp.* (*)** | 9.5 | 19.5 | 29.2 | 31.5 | 54.5 | 91.8 | 122.4 | 31.3 | 2.8 | 24.6 | 4.8 | 1.8 | colony 6 | 1.81 |
| ***Poulsenia armata* (*)** | 32.0 | 13.9 | 31.9 | 14.9 | 55.8 | 25.1 | 104.0 | 31.4 | 11.0 | 19.9 | 4.1 | 2.7 | colony 4 | 2.91 |
| ***Rinorea hummelii* (*)** | 14.0 | 24.2 | 48.7 | 20.7 | 81.2 | 38.2 | 161.5 | 28.0 | 3.4 | 24.8 | 2.9 | 4.2 | colony 2 | 0.02 |
| ***Rinorea hummelii* (*)** | 15.7 | 17.6 | 51.1 | 11.2 | 67.5 | 31.6 | 69.8 | 19.2 | 5.3 | 22.4 | 3.4 | 2.5 | colony 2 | 0.42 |
| ***Serjania mexicana* (*)** | 15.3 | 23.1 | 83.4 | 18.0 | 137.2 | 62.1 | 138.0 | 34.8 | 5.2 | 24.4 | 3.1 | 3.4 | colony 3 | 0.07 |
| ***Swartzia guatemalensis* (*)** | 8.7 | 31.0 | 41.4 | 10.9 | 72.4 | 18.6 | 68.7 | 22.6 | 1.1 | 12.4 | 1.6 | 2.6 | colony 2 | 2.52 |
| ***Swartzia guatemalensis* (***)** | 19.9 | 22.2 | 54.2 | 8.8 | 91.5 | 20.8 | 52.6 | 24.3 | 3.2 | 11.9 | 2.0 | 3.1 | colony 2 | 0.30 |
| ***Swietenia macrophylla* (*)** | 14.4 | 14.9 | 51.2 | 9.8 | 124.3 | 22.2 | 286.5 | 20.3 | 4.3 | 15.6 | 3.3 | 2.6 | colony 4 | 1.52 |
| ***Swietenia macrophylla* (*)** | 18.3 | 20.6 | 47.8 | 9.2 | 56.4 | 16.3 | 46.9 | 15.4 | 1.6 | 9.9 | 1.0 | 1.4 | colony 3 | 0.10 |
| ***Terminalia hainanensis* (*)** | 14.9 | 10.4 | 30.5 | 10.1 | 67.9 | 147.9 | 719.1 | 25.9 | 10.2 | 11.4 | 4.5 | 1.5 | colony 3 | 9.72 |
| ***Terminalia hainanensis* (*)** | 5.7 | 10.2 | 65.6 | 8.3 | 80.2 | 100.2 | 336.8 | 15.8 | 7.9 | 9.5 | 2.6 | 1.5 | colony 3 | 1.52 |
| ***Terminalia hainanensis* (*)** | 14.5 | 11.1 | 19.0 | 9.7 | 19.3 | 77.1 | 212.9 | 21.5 | 7.4 | 12.3 | 2.3 | 1.6 | colony 6 | 0.07 |
| ***Terminalia pallida* (*)** | 9.9 | 12.6 | 42.0 | 14.1 | 67.8 | 33.2 | 161.9 | 23.1 | 5.1 | 15.1 | 2.0 | 2.2 | colony 2 | 1.64 |
| ***Trophis racemosa* (*)** | 31.3 | 21.2 | 137.3 | 9.2 | 195.8 | 91.4 | 41.9 | 42.1 | 4.8 | 17.9 | 2.9 | 3.1 | colony 3 | 14.03 |

| | | | | | | | | | | | | | |
|---|---|---|---|---|---|---|---|---|---|---|---|---|---|
| ***Trophis racemosa* (*)** | 13.1 | 21.3 | 59.9 | 8.9 | 161.5 | 40.2 | 43.7 | 37.3 | 6.4 | 20.5 | 3.8 | 3.8 | colony 3 | 10.05 |
| ***Vitex megapotamica* (*)** | 15.4 | 14.2 | 35.8 | 10.2 | 53.1 | 23.0 | 44.7 | 34.8 | 2.3 | 26.2 | 3.8 | 2.6 | colony 6 | 6.30 |
| ***Vitex megapotamica* (*)** | 14.5 | 25.5 | 64.9 | 28.3 | 116.0 | 211.0 | 116.3 | 84.2 | 2.8 | 38.9 | 6.2 | 4.9 | colony 6 | 0.02 |

For each substrate the table indicates the species identification and substrates type (*: leaves; **: flowers; ***: fruits), the percentages of total non-structural carbohydrates (TNC=water soluble sugars + starch) and protein quantified using NIRS (see methods), the concentration of mineral elements (in ug/g and mg/g) quantify using ICP-OES (see methods), the colony that collected the substrate, and the total dry mass collected. NA indicates non-available data. ǂ refers to negative quantification obtained with NIR predictive model. These values have been considered as NA for statistical analysis.

**Table S9: Intake targets selected by free-ranging colonies of *A. colombica* for carbohydrates, protein and mineral elements.**

| | TNC % | Protein % | Al (ug/g) | Cu (ug/g) | Fe (ug/g) | Mn (ug/g) | Na (ug/g) | Zn (ug/g) | Ca (mg/g) | K (mg/g) | Mg (mg/g) | P (mg/g) |
|---|---|---|---|---|---|---|---|---|---|---|---|---|
| **Colony 1** | 10.38 ± 4.38 | 13.21 ± 1.06 | 41.87 ± 27.0 | 16.08 ± 5.08 | 68.7 ± 35.07 | 144.77 ± 56.5 | 106.63 ± 126.75 | 22.97 ± 3.6 | 5.36 ± 1.24 | 12.52 ± 3.26 | 3.04 ± 0.05 | 2.57 ± 0.92 |
| **Colony 2** | 9.34 ± 1.54 | 18.81 ± 6.5 | 74.91 ± 25.73 | 12.21 ± 1.14 | 92.57 ± 22.36 | 33.40 ± 5.83 | 63.29 ± 25.83 | 29.14 ± 4.73 | 7.37 ± 4.41 | 13.05 ± 1.68 | 2.01 ± 0.23 | 2.25 ± 0.24 |
| **Colony 3** | 18.21 ± 8.6 | 17.65 ± 1.71 | 98.91 ± 25.9 | 9.52 ± 1.31 | 136.7 ± 49.21 | 101.1 ± 58.03 | 319.48 ± 449.8 | 33.14 ± 4.32 | 8.26 ± 3.11 | 16.36 ± 3.53 | 4.12 ± 1.18 | 2.46 ± 0.78 |
| **Colony 4** | 18.74 ± 7.68 | 18.35 ± 4.46 | 211.52 ± 240,7 | 12.73 ± 2.62 | 261.15 ± 293,7 | 37.74 ± 7.62 | 114.03 ± 40.01 | 32.75 ± 9.08 | 7.22 ± 4.21 | 17,76 ± 2.59 | 2,86 ± 0.58 | 2.85 ± 0.99 |
| **Colony 5** | 14.08 ± 8.22 | 17.99 ± 7.09 | 85.07 ± 38.49 | 19.19 ± 9.14 | 108.16 ± 39.26 | 29.02 ± 5.25 | 67.46 ± 13.45 | 37.82 ± 21.54 | 6.16 ± 2.73 | 18.9 ± 5.99 | 2.39 ± 0.35 | 3.54 ± 1.53 |
| **Colony 6** | 13.09 ± 2.04 | 15.58 ± 1.79 | 60.23 ± 40.42 | 13.68 ± 1.1 | 86.47 ± 32.64 | 33.81 ± 10.09 | 72.2 ± 30.06 | 31.93 ± 5.3 | 3.88 ± 0.45 | 17.84 ± 5.27 | 2.78 ± 0.77 | 2.2 ± 0.24 |
| **Mean** | 13.9 ± 3.90 | 16.93 ± 2.14 | 95.4 ±60.21 | 13.9 ± 3.35 | 125.63 ± 70.2 | 63.3 ± 48.28 | 123.85 ± 98.16 | 31.29 ± 4.95 | 7.26 ± 2.97 | 16.07 ± 2.68 | 2,87. ± 0.72 | 2.64 ± 0.50 |

For each colony, values are expressed as the mean of the three weighted means calculated from the three collection days to account for the variation in quantity of plant substrates harvested. The mean (± standard deviation) is calculated for each nutrient from the 6 observed colonies.

### Table S10: Recipes for the synthetic *in vitro* diets used to determine fungal FNNs.

| Nutrient composition of diets | | | | | | |
|---|---|---|---|---|---|---|
| **Pr:C** | **Bacto peptone (g)** | **Bacto tryptone (g)** | **Trypticase peptone (g)** | **Sucrose (g)** | **Starch (g)** | **Agar (g)** |
| **Diet concentration: 4 g/L** | | | | | | |
| 9:1 | 0.064 | 0.064 | 0.064 | 0.010 | 0.010 | 0.800 |
| 6:1 | 0.062 | 0.061 | 0.061 | 0.014 | 0.014 | 0.800 |
| 3:1 | 0.053 | 0.054 | 0.053 | 0.025 | 0.025 | 0.800 |
| 1:1 | 0.036 | 0.036 | 0.036 | 0.050 | 0.050 | 0.800 |
| 1:3 | 0.018 | 0.018 | 0.018 | 0.075 | 0.075 | 0.800 |
| 1:6 | 0.010 | 0.010 | 0.010 | 0.086 | 0.086 | 0.800 |
| 1:9 | 0.007 | 0.007 | 0.007 | 0.090 | 0.090 | 0.800 |
| **Diet concentration: 8 g/L** | | | | | | |
| 9:1 | 0.129 | 0.128 | 0.128 | 0.019 | 0.019 | 0.800 |
| 6:1 | 0.123 | 0.122 | 0.122 | 0.028 | 0.028 | 0.800 |
| 3:1 | 0.107 | 0.108 | 0.107 | 0.049 | 0.049 | 0.800 |
| 1:1 | 0.071 | 0.072 | 0.071 | 0.100 | 0.100 | 0.800 |
| 1:3 | 0.036 | 0.036 | 0.036 | 0.150 | 0.150 | 0.800 |
| 1:6 | 0.020 | 0.020 | 0.020 | 0.171 | 0.171 | 0.800 |
| 1:9 | 0.014 | 0.014 | 0.014 | 0.180 | 0.180 | 0.800 |
| **Diet concentration: 20 g/L** | | | | | | |
| 6:1 | 0.304 | 0.308 | 0.304 | 0.069 | 0.069 | 0.800 |
| 1:1 | 0.177 | 0.180 | 0.178 | 0.249 | 0.249 | 0.800 |
| 1:6 | 0.051 | 0.051 | 0.051 | 0.428 | 0.428 | 0.800 |
| **Diet concentration: 25 g/L** | | | | | | |
| 9:1 | 0.404 | 0.399 | 0.399 | 0.059 | 0.059 | 0.800 |
| 6:1 | 0.384 | 0.380 | 0.380 | 0.087 | 0.087 | 0.800 |
| 3:1 | 0.333 | 0.337 | 0.333 | 0.154 | 0.154 | 0.800 |
| 1:1 | 0.222 | 0.224 | 0.222 | 0.311 | 0.311 | 0.800 |
| 1:3 | 0.111 | 0.113 | 0.111 | 0.468 | 0.468 | 0.800 |
| 1:6 | 0.063 | 0.064 | 0.063 | 0.535 | 0.535 | 0.800 |
| 1:9 | 0.045 | 0.045 | 0.045 | 0.563 | 0.563 | 0.800 |
| **Mineral concentration of diets** | | | | | | |
| **Mineral concentration in media (mg/L):** | | **Aluminum (mL)** $Al_2(SO_4)_3 H_2O$ (10 g/L) (i.e. Al 1.5 g/L) | **Copper (mL)** $CuSO_4 5H_2O$ (10 g/L) (i.e. Cu 2.6 g/L) | **Iron (mL)** $FeSO_4 7H_2O$ (50 g/L) (i.e. Fe 10 g/L) | **Manganese (mL)** $MnCl_2 4H_2O$ (10 g/L) (i.e. Mn 2.77 g/L) | **Zinc (mL)** $ZnSO_4 7H_2O$ (66 g/L) (i.e. Zn 15 g/L) |
| 0 | | 0 | 0 | 0 | 0 | 0 |
| 15 | | 0.50 | 0.29 | 0.075 | 0.27 | 0.05 |
| 30 | | 1.0 | 0.58 | 0.15 | 0.54 | 0.10 |
| 60 | | 2.0 | 1.15 | 0.30 | 1.10 | 0.20 |
| 120 | | 4.0 | 2.3 | 0.60 | 2.16 | 0.40 |
| 240 | | 8.0 | 4.62 | 1.20 | 4.33 | 0.80 |
| 480 | | 16.0 | 9.23 | 2.40 | 8.66 | 1.60 |
| 950 | | 31.7 | 18.3 | 4.75 | 17.15 | 3.20 |
| 1200 | | 40.0 | 23.1 | 6.0 | 21.66 | 4.0 |
| **Mineral concentration in media (mg/L):** | | **Calcium (mL)** $CaCl_2$ (50 g/L) (i.e. Ca 18 g/L) | **Potassium (mL)** KCl (50 g/L) (i.e. K 26.2 g/L) | **Magnesium (mL)** $MgSO_4$ (75 g/L) (i.e. Mg 15.1 g/L) | **Sodium (mL)** NaCl (50 g/L) (i.e. Na 19.7 g/L) | **Phosphorus (mL)** 85% $H_3PO_4$ (i.e. P 454.5 g/L) |
| 0 | | 0 | 0 | 0 | 0 | 0 |
| 150 | | 0.4 | 0.29 | 0.5 | 0.38 | 0.016 |
| 300 | | 0.8 | 0.57 | 1.0 | 0.76 | 0.032 |
| 600 | | 1.7 | 1.16 | 2.0 | 1.52 | 0.065 |
| 1200 | | 3.3 | 2.29 | 4.0 | 3.05 | 0.131 |
| 2400 | | 6.7 | 4.58 | 7.9 | 6.09 | 0.261 |
| 4800 | | 13.3 | 9.16 | 15.9 | 12.18 | 0.523 |
| 9500 | | 26.3 | 18.13 | 31.5 | 24.11 | 1.035 |
| 12000 | | 33.3 | 22.9 | 39.7 | 30.46 | 1.307 |
| **Mineral solutions were prepared with distilled water and all diets were adjusted to a final volume of 50 mL with distilled water.** | | | | | | |

The top panel shows the recipes for the 21 diets varying in Pr:C ratio (9:1, 6:1, 3:1, 1:1, 1:3, 1:6, 1:9) and Pr+C dilutions (4, 8 and 25 g/L). The bottom panel shows the quantity of mineral solutions used.

**Table S11: Descriptive statistics of protein and total non-structural carbohydrate (TNC) contents in plant material for calibration and predicted samples.**

| **Protein %** (PLS regression model → RMSEE: 2.08 / RMSECV: 3.23 / $R^2$: 0.93) | | | | | |
|---|---|---|---|---|---|
| | n | Minimum | Maximum | Mean | SD |
| Calibration | 30 | 2.39 | 31.05 | 14.32 | 7.45 |
| Predicted | 57 | -1.05 | 34.83 | 15.49 | 7.33 |
| **Total non-structural carbohydrate %** (PLS regression model → RMSEE: 9.19 / RMSECV: 10.11 / $R^2$: 0.44) | | | | | |
| | n | Minimum | Maximum | Mean | SD |
| Calibration | 30 | 2.99 | 45.17 | 16.28 | 12.05 |
| Predicted | 57 | 8.02 | 38.35 | 16.59 | 5.44 |

The table shows minimum and maximum values, as well as mean values and the standard deviation for both calibration and predicted samples obtained using wet chemistry and the 1st derivative of the NIR spectra for protein and carbohydrates. n indicates the number of samples for both calibration and predicted dataset. RMSEE, RMSECV and $R^2$ are indicators of quality of the prediction model. RMSEE indicates root mean square error of the estimation for observations in the workset. RMSECV indicates root mean square error computed from the selected cross validation round. $R^2$ indicates the relationship between the measured and predicted samples. Calibration model predicted protein with high prediction accuracy ($R^2$ = 0.93) and TNC with moderate accuracy ($R^2$ = 0.44).

**Legends for Dataset1 (separate file)**.

The data contained in each sheet have been used to generate the figures and tables present in the corresponding sheet title.

Sheet1: "Figures 2, 4C-D, S1, S7, S8" contains mean values for fungus grown on baseline diets with no minerals.

Variables are: Diet (Pr+C concentration and Pr:C ratio of the diets), FinalArea (Hyphal growth in mm2), StaphylaNb (Staphyla number), StaphylaDensity (Staphyla density), Survival (in %), prot (concentration of protein) and carb (concentration of carbohydrates) in diets (both in g/L), pctprot in % (percent of protein) and pctcarb in % (carbohydrates) of diets.

Sheet2: "Tables S1-S2" contains raw data for fungus grown on baseline diets with no minerals.

Variables are: Photo_ID and Plate_ID (identification of plates), Biomass (Pr+C concentration in g/L of the diets), PC (Pr:C ratio of the diets), FinalArea (Hyphal growth in mm2), staphyla_nb (Staphyla number), survival (in %), staph_density (Staphyla density), Biomass_bis (Pr+C concentration changed to factors), prot (concentration of protein) and carb (concentration of carbohydrates) in diets (both in g/L).

Sheet 3 to 12 contains hyphal growth mean data (in mm2) in presence of mineral elements (baseline + 8 mineral concentrations) across Pr:C diets at Pr+C 8 g/L, formated and scaled to 1250 mm2 (the maximum value across all treatments) to build heatmaps mapping hyphal growth.

Sheet3: "Figure S2 Al" contains Al data.
Sheet4: "Figure S2 Ca" contains Ca data.
Sheet5: "Figure S2 Cu" contains Cu data.
Sheet6: "Figure S2 Fe" contains Fe data.
Sheet7: "Figure S2 K" contains K data.
Sheet8: "Figure S2 Mg" contains Mg data.
Sheet9: "Figure S2 Mn" contains Mn data.
Sheet10: "Figure S2 Na" contains Na data.
Sheet11: "Figure S2 P" contains P data.
Sheet12: "Figure S2 Zn" contains Zn data.

Sheet13: "Figure 3, Figure S3" contains data of hyphal growth on diets containing mineral elements. For each mineral is indicated the ID for both concentrations tested (i.e. Al3 and Al5 for aluminum), the final hyphal growth in mm2 (Al3FinalArea) and the percent of growth relative to control in % (Al3ratiopct). This classification applies for all minerals tested.

Variables also contain: Diet (Pr+C concentration and Pr:C ratio of the diets), prot (concentration of protein) and carb (concentration of carbohydrates) in diets (both in g/L), pctprot in % (percent of protein) and pctcarb in % (carbohydrates) of diets.

Sheet14: "Tables S3-S5" contains data (from 3 mineral concentrations (no mineral, best growth, last growth) used for statistical analysis to infer the role of nutrients and mineral elements on hyphal growth.

Variables are: Photo_ID (identification of plates), Biomass (Pr+C concentration in g/L of the diets), PC (Pr:C ratio of the diets), Mineral (Mineral used; C0 means baseline with no mineral), Mineral_concentration (factors indicating the mineral concentrations used), FinalArea (final hyphal growth in mm2), concentration_bis (mineral concentrations in g/L), Biomass_bis (Pr+C concentration changed to factors), prot (concentration of protein) and carb (concentration of carbohydrates) in diets (both in g/L).

Sheet15: "Figures 4, 5A, S5-S9" contains nutrient quantifications for the 87 plant substrates.

Variables are: Species name (Species), Sample_ID (sample identification), TNC (total non-structural carbohydrates in %), Protein (in %), DryMass (in mg), Al (Aluminum concentration in ug/g), Ca (Calcium in ug/g), Cu (Copper in ug/g), Fe (Iron in ug/g), K (Potassium in ug/g), Mg (Magnesium in ug/g), Mn (Manganese in ug/g), Na (Sodium in ug/g), P (Phosphorus in ug/g), Zn (Zinc in ug/g), Colony_ID (identity of the colony that harvested the substrate), Collection_day (Day of collection), Location (Sampling location), Substrate types (either leaves, fruit or flower).

Sheet16: "Figure 4A" contains dry quantity of substrates (quantity in g) for each plant species (species).

Sheet17: "Figure S4" contains the numbers of plant fragments harvested by ants (Harvested_fragments) and collected by samplers (Sampled_fragments) per 30-minute periods for each colony (Colony_ID) for each sampling day (Day).

Sheet18: "Figure S5A" is the matrix to build Venn diagram of plant distribution across locations. 1 indicates presence and 0 indicates abscence of plant species in PLR (Pipeline road), LF (Laguna forest) and CC (Camino de cruces).

Sheet19: "Figure S5B" is the matrix to build Venn diagram of plant distribution across colonies. 1 indicates presence and 0 indicates abscence of plant species for the different colonies (Col1 to Col6).

Sheet20: "Figure 5C" contains mean nutrient quantifications for the different plant samples per species and substrate types.

Variables are: Species name (Species), Type of substrate (Substrate_type), TNCpct (total non-structural carbohydrates in %), Proteinpct (in %), Alconc (Aluminum concentration in µg/g), Caconc (Calcium concentration in µg/g), Cuconc (Copper concentration in µg/g), Feconc (Iron concentration in µg/g), Kconc (Potassium concentration in µg/g), Mgconc (Magnesium concentration in µg/g), Mnconc (Manganese concentration in µg/g), Naconc (Sodium concentration in µg/g), Pconc (Phosphorus concentration in µg/g), Znconc (Zinc concentration in µg/g).

Sheet21: "All data 7PC, 3P+C, 9 min.Conc." contains all raw data of hyphal growth from diets with mineral elements (raw data used to generate mean values in Sheet 9 to Sheet 19, and raw data from Sheet 20).
